# Cumulative effects of high temperature and low dissolved oxygen alter the acute thermal tolerance and cellular stress response in lake trout

**DOI:** 10.1101/2025.02.07.637131

**Authors:** Alyssa M Weinrauch, Analisa Lazaro-Côté, Travis C Durhack, Eva C Enders, Ken M Jeffries

## Abstract

Lake trout (*Salvelinus namaycush*) is an important food fish in Northern communities, inhabiting cool, well-oxygenated water. Yet, climate change is reducing available habitat with extended summer stratification of lakes creating an upper thermal barrier (∼15 °C) and lower dissolved oxygen (DO) boundary (4-7 mg L^-1^). Together, these environmental factors can influence tolerance thresholds and climate change may lead to abiotic factors exceeding these physiological thresholds in lake trout habitats. Thresholds can shift with environmental acclimation in lake trout populations, but the functional basis of this shift has yet to be examined. The abundance of mRNA transcripts offers insight into underlying cellular responses to environmental stressors that can provide an early warning of fitness consequences. Here, we used a stress-response transcriptional profiling chip to investigate a suite of genes involved in thermal and general stress in lake trout acclimated to a range of temperatures (6-18 °C) and DO (normoxia: > 8.5 mg L^-1^ or hypoxia: 5.5-6.5 mg L^-1^), as well as following an acute thermal stress (i.e., CT_max_). Transcriptional profiles were assessed in the gill, liver, and and epidermal mucus. Generally, fish acclimated to the greatest combined stressor (18 °C and hypoxia) had the largest transcriptional response, suggestive of a transition from a routine stress response to an extreme survival response. A noted temperature dependence occurred in liver tissue, which was not evident in gill or mucus tissues. Further, transcriptional responses in the gill and mucus were highly correlated (*r* = 0.74-0.87), highlighting the potential use of these tissues for non-lethal sampling methods to enhance management and conservation strategies for lake trout across their distribution.

## Introduction

Climate change is increasing surface water temperatures and thermal stratification in lakes worldwide, with some of the largest ecological changes occurring at northern high latitudes (Kraemer *et al*., 2021; O’Reilly *et al*., 2015; Woolway *et al*., 2022). Specifically, reduction in ice-cover periods leads to earlier onset and longer duration of thermal stratification, thereby restricting thermal refugia for cold-water inhabitants (Woolway *et al*., 2022). Coupled with higher surface temperatures is a decrease in hypolimnion dissolved oxygen (DO) concentration (Jane *et al*., 2024; Jansen *et al*., 2024). This occurs as a result of reduced atmospheric oxygen exchange during extended stratification and increased dissolved organic carbon loading leading to increased microbial respiration in warmer waters (Jane *et al*., 2024; Jansen *et al*., 2024). Thus, cold-water animals face a reduced ecological niche (i.e., habitat ‘squeeze’), wherein only a narrow region of the water column is suitable habitat during parts of the year (Lyons *et al*., 2018; Woolway *et al*., 2022). Together, these oxythermal habitat alterations will affect northern high latitude lake inhabitants to a varying degree depending upon their oxythermal tolerance, with greater effects predicted for cold-water ectotherms (Kraemer *et al*., 2021).

Lake trout (*Salvelinus namaycush*) is a cold-water fish, native to temperate and subarctic lakes across the boreal shield in Canada (Scott and Crossman 1973). It is an important subsistence food fish for many Northern and Indigenous communities (Kuhnlein & Humphries, 2017). As a top predator with a preferred habitat of cool (8-12 °C), well-oxygenated water (lower boundary 4-7 mg L^-1^; Blanchfield et al., 2009; Christie and Regier, 1988; Clark et al., 2004; Evans, 2007; Martin and Olver, 1980), lake trout play a key role in nutrient cycling within lake ecosystems (Ivanova *et al*., 2021). Yet, being ectotherms, their distribution is largely governed by temperature (Blanchfield *et al*., 2009), and climate warming can thus have profound effects on lake ecosystem dynamics via altered fish distributions, which in turn can affect Indigenous communities both in terms of food security and culture. Further, lake trout themselves face physiological consequences when exposed to suboptimal environments. For example, warmer summers correlate with reduced growth and body condition of lake trout owing to a change in foraging from larger littoral prey to smaller pelagic prey that reside within their preferred oxythermal habitat (Guzzo *et al*., 2017). With climate-driven reductions in suitable habitat, lake trout will increasingly face environments that surpass their oxythermal thresholds, necessitating plasticity to cope with these perturbations (Schulte *et al*., 2011). Such plasticity has been shown across lake trout populations where thermal tolerance improved with acclimation to elevated temperature (Kelly *et al*., 2014). Yet, the influence of combined oxygen and thermal stressors on the underlying cellular responses that drive plasticity of thermal thresholds remain unstudied.

Transcriptional profiles (i.e., mRNA abundance) are valuable markers for evidence of plasticity (Akbarzadeh *et al*., 2018, 2020; Jeffries *et al*., 2021; Swirplies *et al*., 2019). Further, these profiles can be used to characterize sublethal cellular responses to environmental perturbation with thresholds ranging from an adaptive response to routine increases in stressors to an extreme stress response characterized by high expression of inducible heat shock proteins, oxidative stress, and cell death mechanisms (reviewed in Jeffries *et al*., 2018). In combination with measures of thermal tolerance (e.g., critical thermal maximum [CT_max_]), sublethal transcriptional profiles can be a useful predictive management tool to identify environmental conditions in which wild fish populations are adversely affected, determine what their thermal limits are, and subsequently provide an early warning of fitness consequences (Akbarzadeh *et al*., 2018, 2020; Jeffries *et al*., 2018; Swirplies *et al*., 2019). Recent studies used a stress-response transcriptional profiling (STP) chip to understand the molecular underpinnings of stress tolerance in salmonids (Best et al., 2024; Islam et al., 2024). These “chips” are a high-throughput qPCR approach that uses OpenArray^®^ technology (Thermo Scientific) to simultaneously assess transcriptional abundance of 52 genes of interest (and four reference genes) across eight diverse functional classifications including: apoptosis, circadian rhythm, detoxification, growth and metabolism, hypoxia, immune function, osmoregulation, and stress response (SI Table 1). Quantification of genes across this diverse array of functional classifications enables physiological interpretation of the cellular stress response, particularly when profiles are examined across multiple tissue types (Semeniuk *et al*., 2022), including epidermal mucus (Andrzejczyk *et al*., 2022).

**Table 1.** Overall average temperature and dissolved oxygen (DO) concentrations in each treatment tank.

| Treatment | Temperature (°C)<br>(mean ± sd) | DO (mg/L)<br>(mean ± sd) |
| --- | --- | --- |
| 6 °C normoxia | 5.82 ± 0.04 | 11.60 ± 0.28 |
| 10 °C normoxia | 9.82 ± 0.04 | 10.19 ± 0.68 |
| 14 °C normoxia | 13.86 ± 0.07 | 9.38 ± 0.66 |
| 18 °C normoxia | 18.04 ± 0.04 | 8.67 ± 0.52 |
| 6 °C hypoxia | 5.85 ± 0.16 | 6.18 ± 0.44 |
| 10 °C hypoxia | 10.08 ± 0.76 | 6.17 ± 0.33 |
| 14 °C hypoxia | 13.71 ± 0.83 | 6.17 ± 0.56 |
| 18 °C hypoxia | 18.2 ± 0.18 | 5.37 ± 0.72 |
| Normoxia_2 at 10 °C | 9.94 ± 0.91 | 10.06 ± 0.37 |

The purpose of this study was to examine the cellular stress response of the stenothermal lake trout subjected to ecologically-relevant oxythermal stressors. Specifically, fish were acclimated to a range of temperatures (6, 10, 14, 18 °C) and two DO concentrations (9.93 ± 1.23 mg L^-1^ hereafter referred to as normoxia and a reduced DO treatment of 6.18 ± 0.44 mg L^-1^ hereafter referred to as hypoxia) that encompass ecologically relevant water temperature and oxygen levels including and beyond preferred habitat. Further, the transcriptional profile was assessed in individuals that underwent an acute thermal stress (i.e., CT_max_), to test for plasticity in thermal tolerance and its molecular underpinnings. We predicted that the cellular stress response would shift from a routine stress response to an extreme stress response in acclimation temperatures above the preferred 10 °C, with a greater shift occurring after CT_max_. We assessed this transcriptional profile in three different tissues (i.e., liver, gill, and epidermal mucus) to permit a more comprehensive interpretation of physiological responses and to shed light on which tissues share a conserved response; thereby identifying non-lethal targets useful for management and conservation of wild lake trout populations.

## Materials and Methods

### Animals and Permits

Lake trout gametes were collected from parental fish in four lakes at the IISD-Experimental Lakes Area (Lakes 260, 223, 375, 378) in the fall of 2019 and brought to the Freshwater Institute in Winnipeg, MB, Canada for fertilization and hatching. Following swim-up, fish were maintained in 200 and 800 L tanks with flow-through de-chlorinated City of Winnipeg tap water with supplemental aeration to maintain dissolved oxygen levels above 80%. Fish were held at 10 °C ± 1 °C on a 12-12 photoperiod and fed a daily ration of 1% total tank mass of commercial food pellets (EWOS Pacific: Complete Fish Feed for Salmonids, Cargill, MN, USA) prior to experimentation. All fish collection and experiments complied with the Canadian Council for Animal Care and were conducted with approval under permits from the Ontario Ministry of Natural Resources (license no. 1094448), University of Manitoba Animal Care Committee (Animal Use Protocol F18-018/1), and Fisheries and Oceans Canada Ontario, Prairie, and Arctic Animal Care Committee (Animal Use Protocols FWI-ACC AUP 2021-72 and OPA-ACC AUP 2022-02).

### Experimental Acclimations and Sampling

The experimental treatments included four temperatures (i.e., 6, 10, 14, 18 °C) and two oxygen concentrations (i.e., normoxia DO = 9.93 ± 1.23 mg L^-1^; hypoxia DO = 6.18 ± 0.44 mg L^-1^; Table 1). Fish (*n* = 25) were indiscriminately selected from the general population and placed into each 200 L treatment tank, with *n* = 10 fish used as controls (i.e., acclimated) and *n* = 15 as acutely heat stressed (i.e., CT_max_) fish. After 24 h of habituation to the treatment tanks, the water was gradually warmed or cooled (2 °C per day) to the appropriate temperature as recorded by a Witrox-4 Instrument and AutoResp Software (Loligo^®^ Systems, Viborg, Denmark). Lake trout were maintained in their respective treatment groups for a period of three weeks during which oxygen concentration was maintained above 90% saturation in normoxia treatments or reduced to ∼ 6 mg L^-1^ in the hypoxia group by controlling aeration via a 1-channel oxygen analyzer (Loligo^®^ Systems, Viborg, Denmark). Of note, owing to constraints of space, both oxygen concentrations could not be conducted simultaneously. Thus, normoxia trials were performed from Dec 7-10, 2021, and hypoxia trials were performed from Jan 24-29, 2022. To ensure that any detected differences were not the result of time alone, a second normoxia trial at 10 °C (referred to as “normoxia_2”) was conducted alongside with the hypoxia trials and compared against the original 10 °C normoxia trial. Responses to treatment across times were virtually identical (see supplement), therefore, data from normoxia_2 are not presented.

At the end of the acclimation period, *n* = 10 lake trout were euthanized with tricaine methanesulfonate (MS222 450 mg L^-1^; Syndel Canada, Nanaimo, BC, Canada) buffered with sodium bicarbonate (900 mg L^-1^; Fisher Scientific, Ottawa, ON, Canada), and measures of fork length and body mass were recorded. Epidermal mucus was collected by rubbing sterile swabs (Mantacc 96000A 6” flocked sampling swab; Miraclean Technology, Guangdong, P.R. China) across the dorsal surface of the right side of the fish (i.e., midline and up). Unfortunately due to a lack of supply, swabs were not collected for the 18 °C normoxia CT_max_ trial. Samples were placed in RNAse-free 1.5 mL microfuge tubes and immediately frozen under liquid nitrogen and stored at -80 °C until further processing. Gill tissue from the second gill arch were collected. The liver was excised, weighed, and a portion of the distal lobe was collected. Both tissues were placed into RNAlater (Thermo Fisher Scientific, Waltham, MA, USA) and stored at -80 °C.

### CT_max_ Experiments

Following the above acclimation protocol, fish (*n =* 15 per treatment) underwent a CT_max_ experiment as described in (Morrison *et al*., 2020). Treatment tanks were heated at a rate of 0.3 °C min^-1^ using six 300 W titanium heaters (Finnex TH-0300S titanium heaters, Finnex, Chicago, IL, USA). Heating rate was recorded using a Witrox-4 instrument and AutoResp software and monitored visually using a TMP-Reg instrument (Loligo^®^ Systems, Viborg, Denmark). Desired dissolved oxygen concentraations were maintained using aerators (and the 1-channel oxygen analyzer for hypoxia treatments) and homogenous temperature throughout the tank was maintained using two 5 L min^-1^ pumps (Eheim, Deizisau, Germany). Fish were constantly monitored by two observers during trials and CT_max_ for individual fish was determined as when loss of equilibrium (LOE) occurred. Following LOE, fish were removed from the tank and euthranized as described above for tissue sampling.

### Measurement of Transcript Abundance

Transcript abundance was quantified using custom designed OpenArray^®^ high-throughput qPCR chips (Thermo Fisher Scientific(Islam *et al*., 2024). These chips accommodated 24 samples run in duplicate (i.e., 2688 simultaneous reactions) for a total of 56 genes (SI Table 1). Briefly, total RNA was extracted using the MagMAX *mir*Vana Total RNA Isolation Kit (Applied Biosystems) with the KingFisher Duo Prime (Thermo Fisher Scientific) according to the manufacturer’s specifications with the following modifications. Mucus swabs were combined with 300 µL of lysis buffer while all other tissues were combined with 600 µL, prior to homogenization (30 s for mucus, 5 min for liver and gill) with a bead homogenizer. Gill and liver samples were eluted in 50 µL of elution buffer while mucus samples were eluted in 25 µL. RNA integrity and purity was assessed using a Nanodrop One. Genomic DNA was removed using ezDNase (Invitrogen) prior to cDNA synthesis (1 µg for gill and liver, 0.25 µg for mucus) with a High Capacity cDNA Reverse Transcription Kit (Applied Biosystems) as per the manufacturer’s protocol. Duplicate samples were diluted 1:4 with nuclease free water and 2.5 µL was loaded alongside 2.5 µL of 2X TaqMan OpenArray Real-Time PCR Master Mix onto the custom designed salmonid stress chips (Best et al. 2024) using the QuantStudio 12K Flex OpenArray^®^ Accufill System (Applied Biosystems). Within 90 s of loading, lids were attached to the chips which were then filled with oil and sealed as per the manufacturer’s specifications. Chips were then run on a QuantStudio^TM^ 12K Flex Real-Time PCR System (Applied Biosystems). Post-run images were analyzed to ensure proper sample loading, and samples where there were loading issues were removed from the analysis.

### Data and Stastical Analyses

All analyses and graphing was conducted in Rstudio v.4.3.2 (https://www.r-project.org/). For CT_max_, a Wilcoxon rank sum test was conducted between the temporally distinct normoxia 10 °C treatments. Finding no significant difference, the second normoxia 10 °C treatment was removed from further analysis. A linear model was implemented in the ‘aov’ function of the *stats* package to investigate effects of temperature acclimation, oxygen concentration, and their interaction on CT_max_. Assumptions of the model were visually inspected with the ‘check_model’ function of the *performance* package v. 0.10.8 (Lüdecke *et al*., 2021). Contrasts of estimated marginal means were computed with the package *emmeans* v.1.9.0 (Lenth, 2021).

For gene transcript analysis, the general protocols of Best et al. (2024) were followed. Briefly, QuantStudio Design and Analysis v1.5.2 and ExpressionSuite software v1.3 (Thermo Fisher Scientific) generated raw amplification data which was screened for quality. Samples that were removed at this stage included no template controls as well as samples that did not amplify, with an amplification score threshold < 1.24, cq confidence threshold < 0.8, or a standard deviation threshold < 0.7. Further, genes with > 50% amplification fail were removed from subsequent analysis. For liver, that included HSP90AA, LDHA, MTB, HIF1α, LEPR, VEGFC, PCK1; for gill: HSP90AA, MTB, HIF1α, CHMP5AB, IGFBP, PCK1; for mucus: GHR, IGF1, HSD11β2, SERPINH1, HSF1, PCK1, HSP90AA, CPT1A, LDHA, MMP2, LPL, IGFBP1, MTB, HIF1α, VEGFC, CRY1AB, CAT, LEPR. Additionally, the PCR efficiency of each amplicon was calculated using LinRegPCR (Ruijter *et al*., 2009). Datasets were then split into control, acclimated fish versus those exposed to a CT_max_ trial. Raw Cq values were converted to counts using the function cq2counts and were analysed with Poisson-lognormal generalized linear models using a Bayesian framework with MCMC.qpcr (doi: 10.32613/CRNA.package.MCMC.qpcr) with MCMC settings of 13,000 iterations, with the first 3,000 iterations discarded, and a thinning interval of parameters sampled once every 10 iterations. Naïve models were first fit to establish whether global effects were evident and if the reference genes (*elf1α*, *rpl7*, *rpl13a*, *rps9*) were stable (see Supplement and SI Fig. 7-12). Upon confirmation of stability, a second model was built using only the stable reference genes as priors and the function diagnostic.mcmc was used to generate diagnostic plots to visually assess model fit. Results were plotted using the package ggplot2 (https://ggplot2.tidyverse.org).

The converted count data generated for MCMC.qpcr was used to create principal component analysis (PCA) plots with the package ‘FactoMineR’ (Lê *et al*., 2008). The top contributing genes to all PCs were established using ‘fviz_cos2’ of the package *factoextra* (10.32614/CRAN.package.factoextra). The normalized counts for each tissue were utilized in a Pearson’s correlation analysis so each tissue type had its own internal control. Correlations were calculated with the function ‘cor’ with plots generated using the R package *corrplot* (v. 0.92).

## Results

Oxygen dependent thermal sensitivity was found depending upon acclimation temperature (Fig. 1). Specifically, differences in upper thermal maximum (CT_max_) were identified with respect to acclimation temperature (F_3,112_ = 127.48, *p* < 0.0001), oxygen concentration (F_1,112_ = 41.03, *p* < 0.0001), and their interaction (F_3,112_ = 7.62, *p* = 0.0001). Specifically, the effect of temperature on CT_max_ showed oxygen dependence, where CT_max_ was significantly higher at normoxia compared to hypoxia except for at 14 °C. The hepatosomatic index (HSI) significantly decreased as temperature increased with a more pronounced decrease in the hypoxia treatment group at 6 and 18 °C observed (SI Table 2; *p* < 0.001).

**Figure 1.**
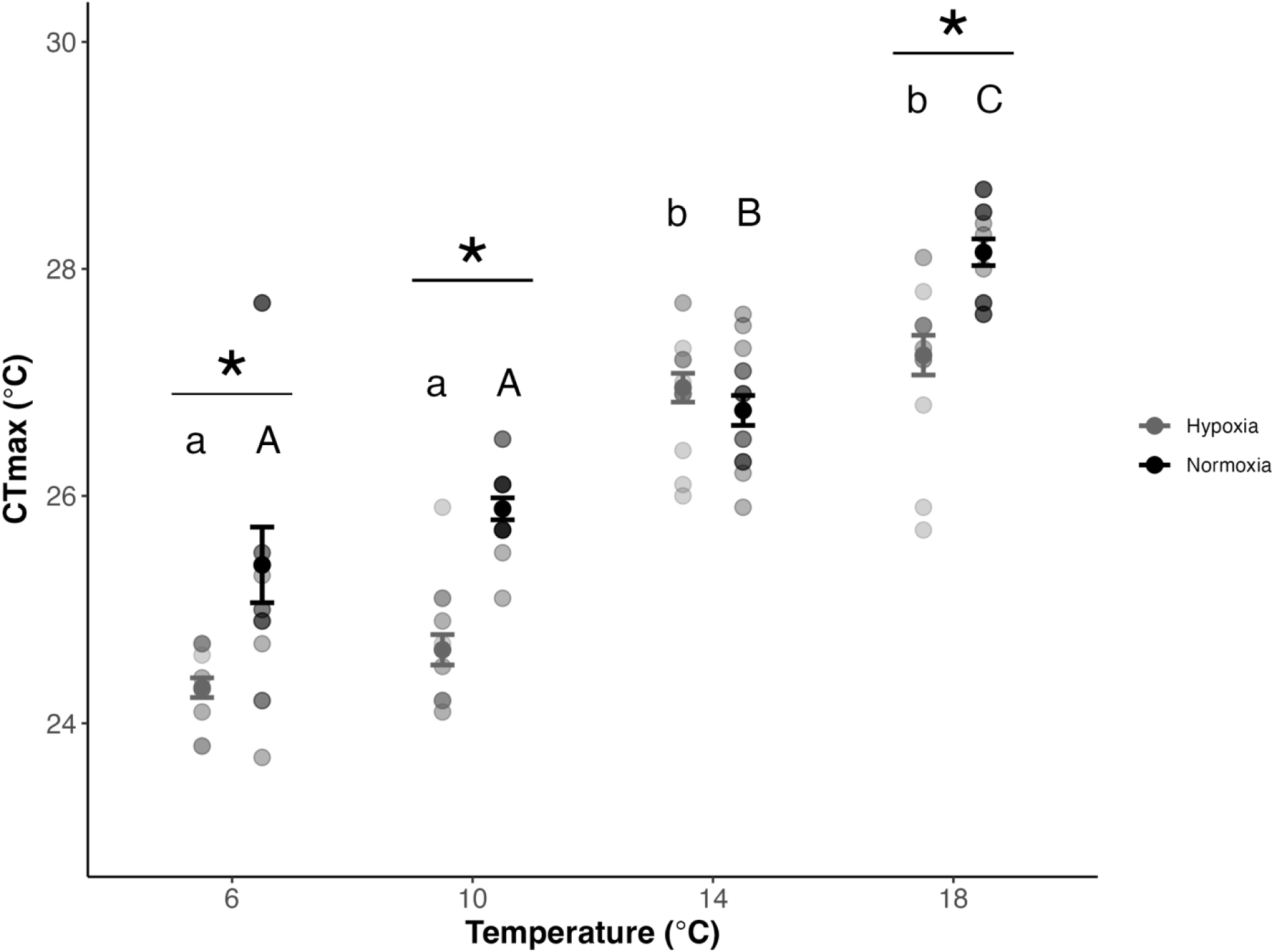
Effects of acclimation temperature and oxygen concentration on thermal tolerance of lake trout (*Salvelinus namaycush*). Lake trout (*n* = 15 per treatment) were acclimated to 6, 10, 14, or 18 °C and normoxia (9.93 ± 1.23 mg L^-1^; black) or hypoxia (6.18 ± 0.44 mg L^-1^; dark grey) for three weeks and critical thermal maximum (CT_max_) was determined at a ramp rate of 0.3 °C min^-1^. Data are presented as means ± s.e.m. with individual data points shown. Asterisks denote significant differences within temperatures while different letters denote significant differences within an oxygen concentration but across temperatures (*p* < 0.05) as determined by a linear model with contrasts estimated using emmeans.

Of the 52 genes of interest on the STP chip, 45 successfully amplified in the liver (SI Fig. 13). A PCA conducted on liver transcript abundance revealed a clustering of samples by temperature (Fig. 2A). Along PC1 (49.6% variance explained), fish were separated primarily by temperature, while PC3 (5.8 % variance explained) showed some separation by dissolved oxygen, with the hypoxia-18 °C fish showing the greatest separation. The top six genes with the greatest joint contribution to PC1 and PC3 were citrate synthase (*cs*; metabolism and growth), caspase 3 (*casp3ab*; apoptosis regulation), superoxide dismutase 1 (*sod1*; oxidative stress protection), sodium-potassium ATPase (*atp1b1*; establishes cellular electrochemical potential), heat shock 70kDa protein a 4 (*hspa4*; stress responsive), and calmodulin (*cam*; mediates numerous processes including immune function). These genes all showed pronounced decreases in transcript abundance with temperature, with some (notably *sod1* and *cam*) demonstrating oxygen-dependent effects at the most extreme temperatures (i.e., 6 and 18 °C; Fig. 2B). Specifically, at 6 °C and 18 °C, the transcript abundance of *sod1* and *cam* was markedly lower for fish held at hypoxia compared to normoxia.

**Figure 2.**
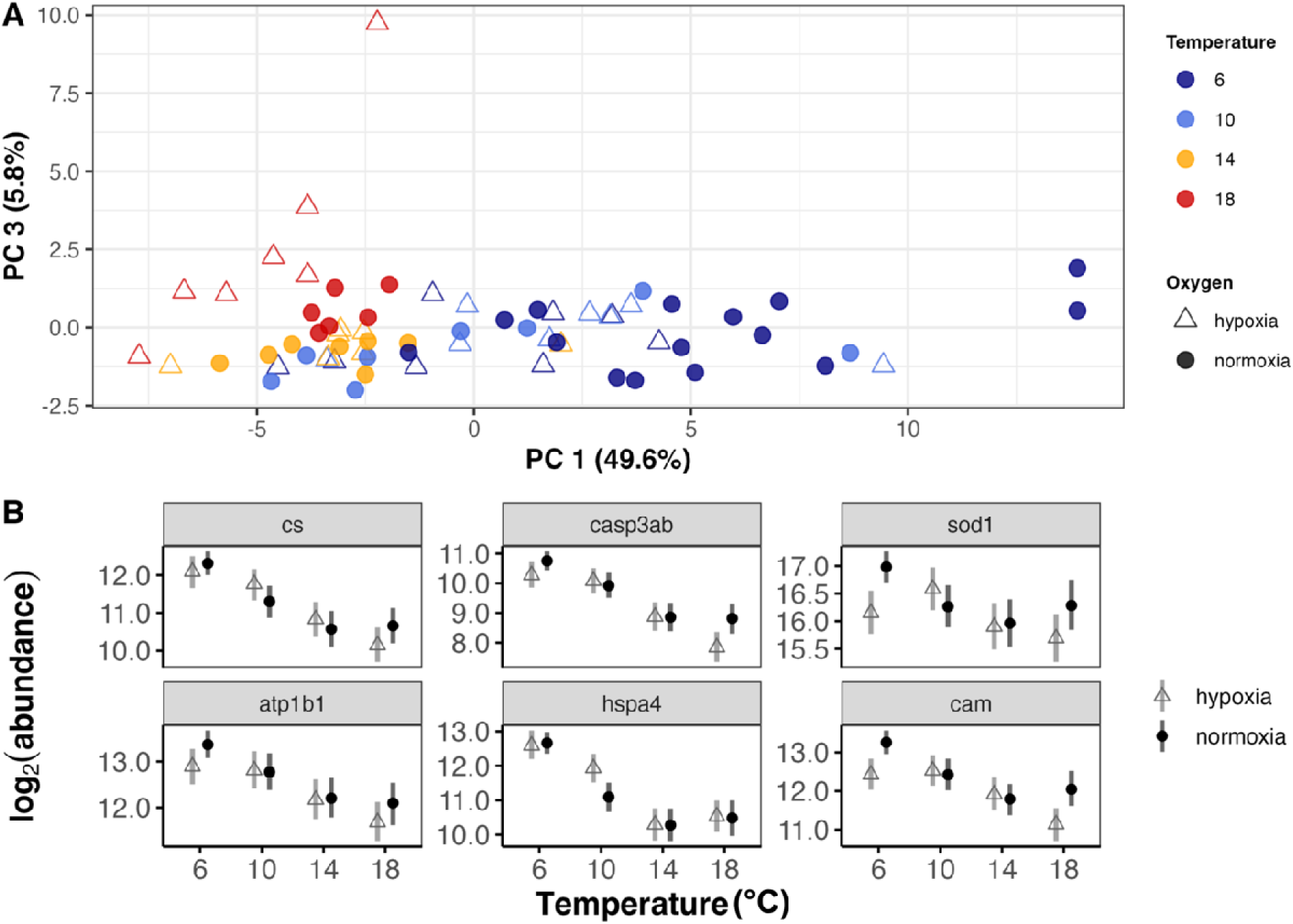
Patterns of mRNA transcript abundance of lake trout (*Salvelinus namaycush*) liver across four acclimation temperatures and two oxygen concentrations. A) Principal components analysis (PCA) of the liver mRNA transcripts for 45 genes of fish (*n* = 6-15 per treatment) acclimated for three weeks to each temperature (colours) and oxygen concentration (shapes). The variance explained by each PC is indicated in brackets. B) Transcript abundance (in log2 abundance) across temperature and normoxia (9.93 ± 1.23 mg L^-1^; black circles) or hypoxia (6.18 ± 0.44 mg L^-1^; grey triangles) for the top six genes contributing to all PC generated in panel A. Data are presented as the posterior means with the whiskers denoting the 95% credible intervals as modeled by the Bayesian analysis tool MCMC.qpcr. Bars that do not overlap are considered significantly different. Note: Models were fit with endogenous control genes (rpl7) as priors. *cs*: citrate synthase, *casp3ab*: caspase 3, *sod1*: superoxide dismutase 1, *atp1b1*: sodium-potassium ATPase, *hspa4*: heat shock protein a4, *cam*: calmodulin.

Forty-six of the target genes successfully amplified in the gill tissue (SI Fig. 14). The first two PCs explained 65.9% of the variation in gene transcript abundance in the gill tissue of control-acclimated fish (Fig. 3A). The variables driving separation on PC1 explained 56.2% of the variance while the variables driving separation on PC2 explained 9.7% of variance. Fish exposed to the highest temperature (i.e., 18 °C) in an hypoxic environment showed the greatest separation from the other treatments. The genes with the highest joint contribution to PC1 and PC2 included superoxide dismutase 2 (*sod2*; detoxification and protection from oxidative stress), *hspa4*, *cam*, *atp1b1*, insulin-like growth factor 2 (*igf2*; growth regulation) and cold inducible RNA binding protein a (*cirbpa*; stress response), three of which were also top contributors in the liver. The genes with the greatest contribution to the overall PCA did not show distinct differences across oxygen concentrations or acclimation temperatures with the exception of *hspa4*, which did have a significant increase in transcript abundance at 18 °C relative to 14 °C (Fig. 3B).

**Figure 3.**
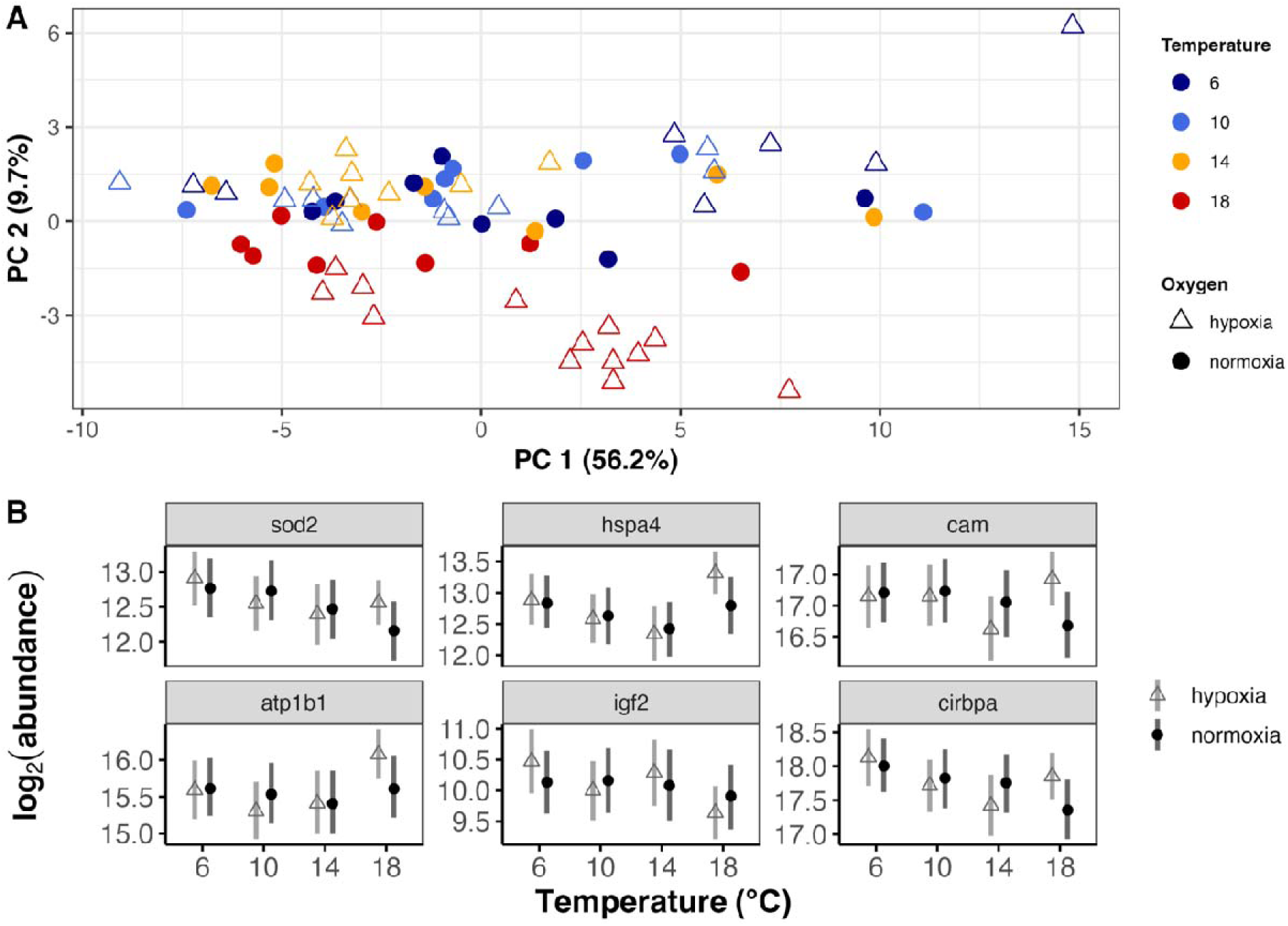
Patterns of mRNA transcript abundance of lake trout (*Salvelinus namaycush*) gill across four acclimation temperatures and two oxygen concentrations. A) Principal components analysis (PCA) of the gill mRNA transcripts for 46 genes of fish (*n* = 8-13) acclimated for three weeks to each temperature (colours) and oxygen concentration (shapes). B) Transcript abundance (in log2 abundance) across temperature and normoxia (9.93 ± 1.23 mg L^-1^; black circles) or hypoxia (6.18 ± 0.44 mg L^-1^; grey triangles) for the top six genes contributing to all PCs generated in panel A. Data are presented as the posterior means with the whiskers denoting the 95% credible intervals as modeled by the Bayesian analysis tool MCMC.qpcr. Bar that do not overlap are considered significantly different. Note: Models were fit with endogenous control genes (*rpl7*, *rpl13*, *rps9*) as priors. *sod2*: superoxide dismutase 2, *hspa4*: heat shock protein a 4, *cam*: calmodulin, *atp1b1*; sodium-potassium ATPase, *igf2*: insulin-like growth factor 2, *cirbpa*: cold inducible RNA binding protein a.

Fewer genes were successfully amplified from epidermal mucus samples (i.e., 32 out of 52; SI Fig. 15). The total variance explained by the first two PCs was 66.2% (Fig. 4A). Interestingly, the 18 °C hypoxia fish did not separate along either principal component using the transcript abundance of genes measured in the mucus like they had in the liver and gill. Examining the six genes with the greatest joint contribution to PC1 and PC2 revealed a consistent pattern for *cirbpa,* aldolase (*aldoaa*; metabolism and hypoxia-responsive), *cam*, *atp1b1,* charged multivesicular body protein 5 (*chmp5ab*; apoptosis) and glutathione S-transferase 1 (*gstp1*; detoxification/oxidative stress response), where transcript abundance was significantly reduced at 10 °C relative to 6 °C. Transcript abundance increased at 14 °C and 18 °C to levels equal to or near 6 °C (Fig. 4B). Finally, *cam* and *gstp1* showed clear separation by oxygen concentration at 10 °C.

**Figure 4.**
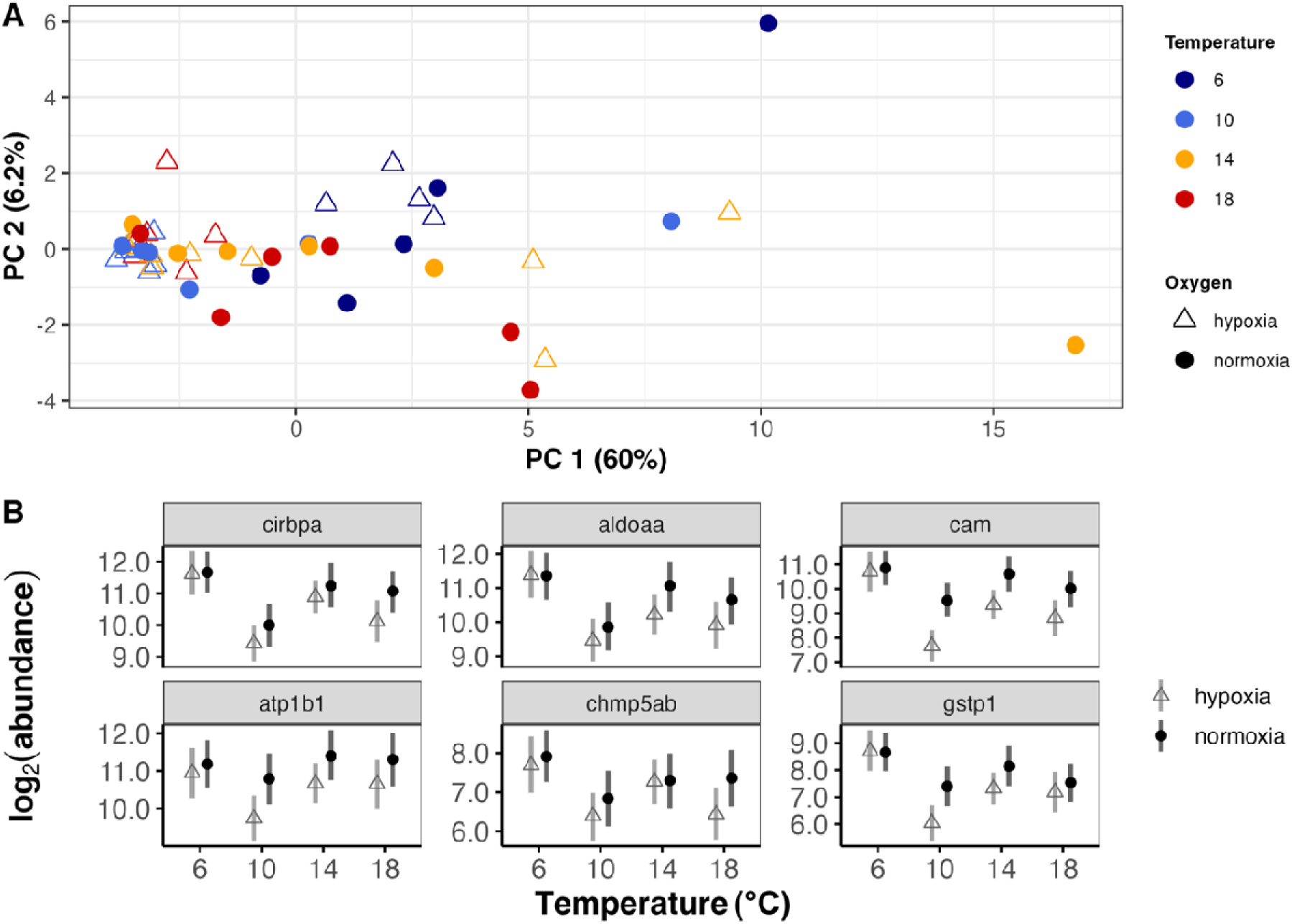
Patterns of mRNA transcript abundance of lake trout (*Salvelinus namaycush*) epidermal mucus swabs across four acclimation temperatures and two oxygen concentrations. A) Principal components analysis (PCA) of the mucus mRNA transcripts for 28 genes of fish (*n* = 4-10) acclimated for three weeks to each temperature (colours) and oxygen concentration (shapes). B) Transcript abundance (in log2 abundance) across temperature and normoxia (9.93 ± 1.23 mg L^-1^; black circles) or hypoxia (6.18 ± 0.44 mg L^-1^; grey triangles) for the top six genes contributing to all PCs generated in panel A. Data are presented as the posterior means with the whiskers denoting the 95% credible intervals as modeled by the Bayesian analysis tool MCMC.qpcr. Bars that do not overlap are considered significantly different. Note: Models were fit with endogenous control genes (*rpl7*, *rpl13*, *rps9*, *ef1α*) as priors. *cirbpa*: cold inducible RNA binding protein a, *aldoaa*: aldolase, *cam*: calmodulin, *atp1b1*: sodium-potassium ATPase, *chmp5ab*: charged multivesicular body protein 5, *gstp1*: glutathione S-transferase 1.

Fish that underwent a CT_max_ trial showed substantial separation of transcriptional profile in the 18 °C fish in both the liver (Fig. 5A) and gill (Fig. 5B). Greater separation was noted across oxythermal regimes for the gill. Similar distinctions at 18 °C between oxygen concentrations were noted for the transcript abundance in both tissues (Fig. 5C, D). The genes comprising the top six contributors to all PCs were different in the CT_max_ fish compared to th control, acclimated fish with the exception of *sod1* in the liver and *igf2* in the gill.

**Figure 5.**
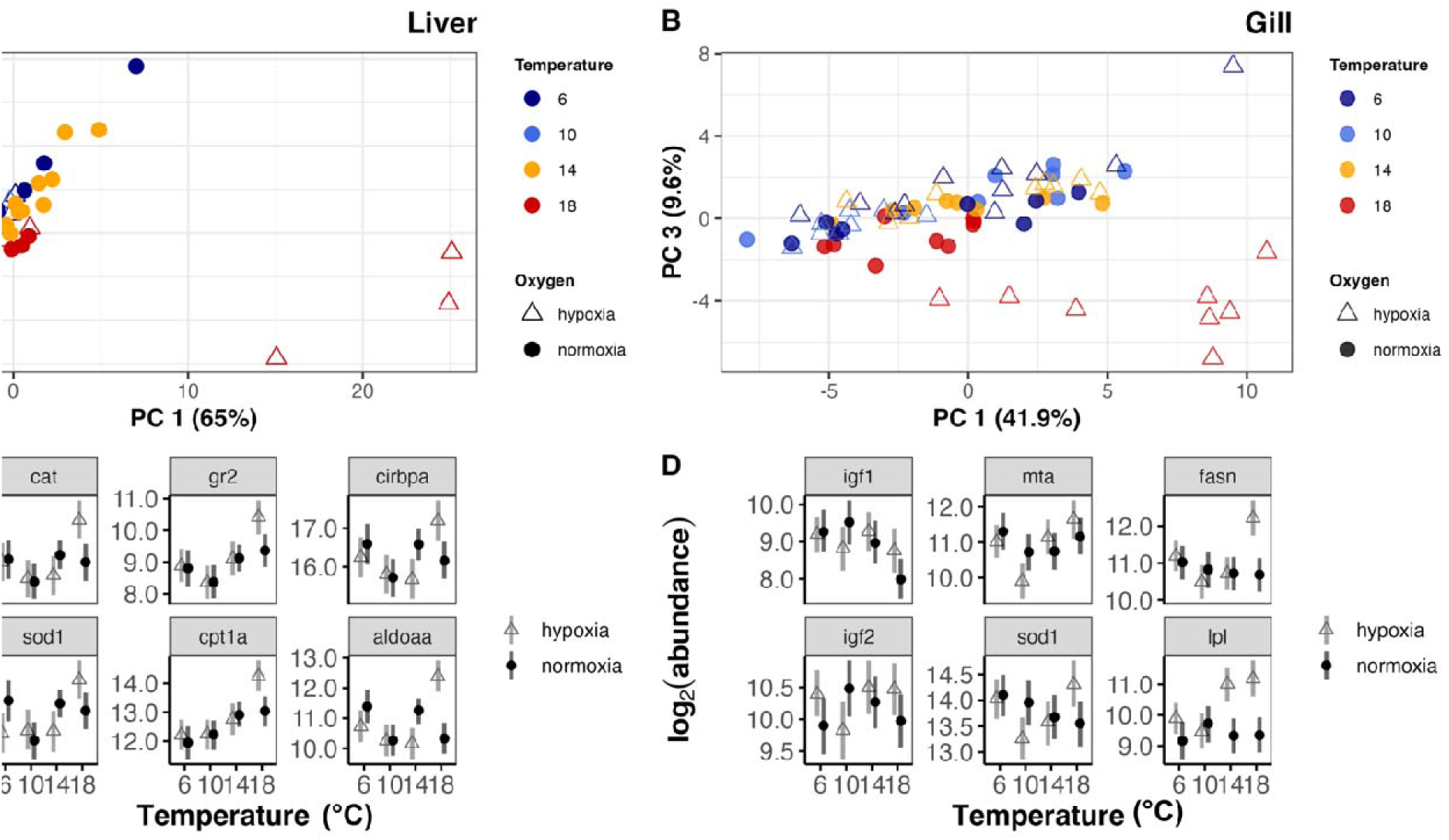
Principal component analysis for mRNA transcript abundance profiles across tissues of lake trout (*Salvelinus namaycush*) following a critical thermal maximum trial. PCAs shown for liver (A; *n* = 6-12) and gill (B; *n* = 8-11) samples from fish acclimated to 6, 10, 14, or 18 °C (colours) and either normoxic (9.93 ± 1.23 mg L^-1^; filled-in circles) or hypoxic (6.18 ± 0.44 mg L^-1^; open triangles) conditions for three weeks. Gene expression patterns modeled by MCMC.qpcr for the top six genes contributing to all PCs for liver (C) and gill (D). Transcript abundance is shown across the four acclimation temperatures for lake trout acclimated to normoxia (black circles) or hypoxia (grey triangles). Abundance data is presented as the posterior means with the whiskers denoting the 95% credible intervals as modeled by the Bayesian analysis tool MCMC.qpcr. Bars that do not overlap are considered significantly different.

The mucus samples showed similar separation as the control, acclimated fish, wherein the 18 °C hypoxia fish did not separate on PC1 or PC2, although a separation by temperature wa evident along PC1 (65% variance explained; Fig. 6A). Mucus samples were not collected at 18°C normoxia, therefore transcript abundance at this temperature is not presented. The top six genes contributing to PC1 and PC2 showed a similar pattern as described for the mucus sample collected from the control acclimated fish wherein transcript abundance decreased at 10 °C, particularly for hypoxia-exposed fish (Fig. 6B).

**Figure 6.**
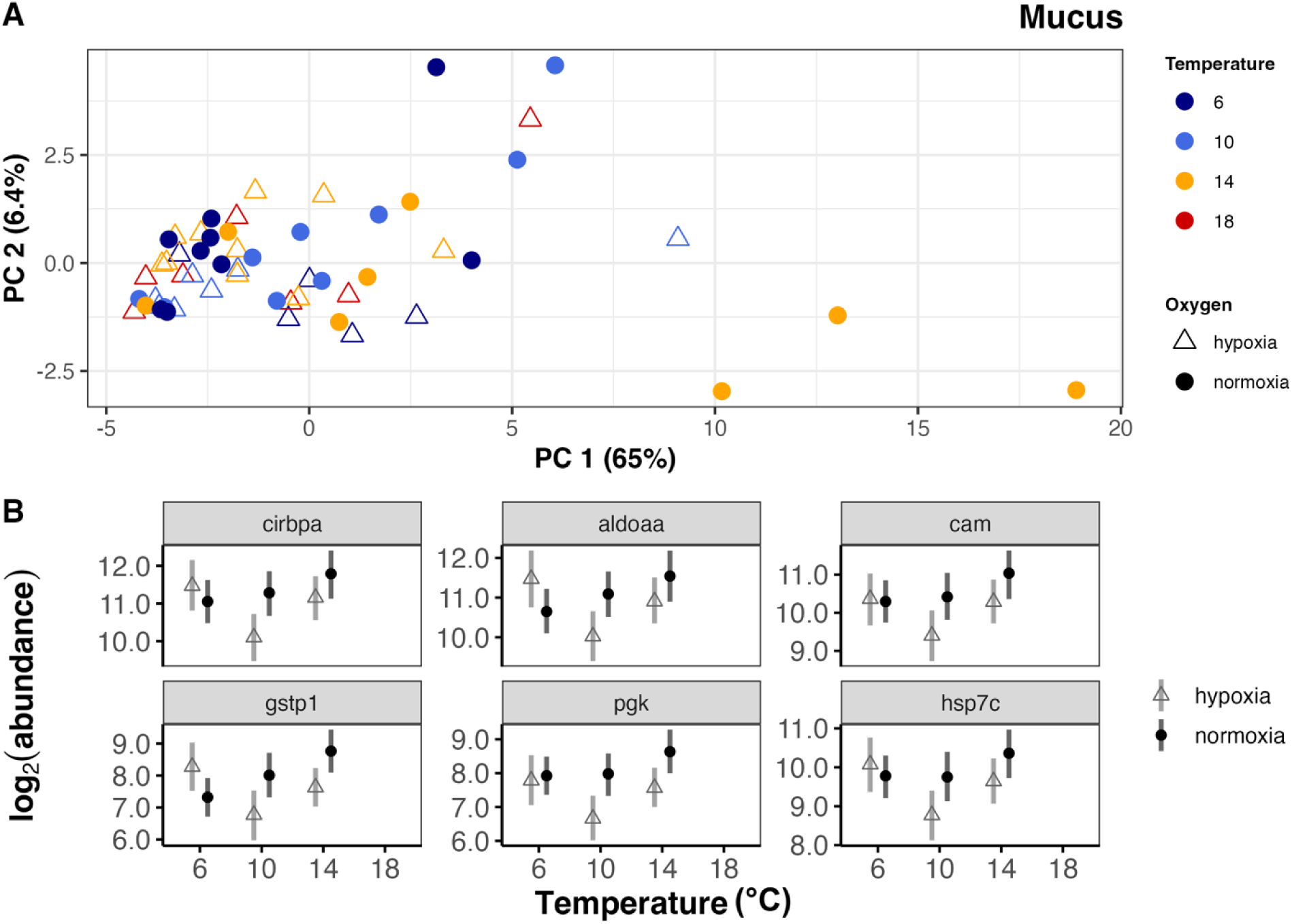
Principal component analysis for gene expression profiles across mucus sample of lake trout (*Salvelinus namaycush*) following a critical thermal maximum trial. A) PCA shown for mucus samples from fish acclimated to 6, 10, 14, or 18 °C (colours) and either normoxic (9.93 ± 1.23 mg L^-1^; filled-in circles) or hypoxic (6.18 ± 0.44 mg L^-1^; open triangles) conditions for three weeks. B) Gene expression patterns modeled by MCMC.qpcr for the top six genes contributing to all PCs (*n* = 4-9). Data are presented as the posterior means with the whiskers denoting the 95% credible intervals as modeled by the Bayesian analysis tool MCMC.qpcr. Bars that do not overlap are considered significantly different.

Correlation analysis demonstrated that transcript abundance in gill and mucus sample were highly correlated across all tested temperatures within both the normoxic and hypoxic groups (*r* = 0.74-0.87; Figure 7). Lower correlation coefficients were observed when comparing the liver and gill (*r* = 0.55-0.60) and the liver and mucus (*r* = 0.49-0.57).

**Figure 7.**
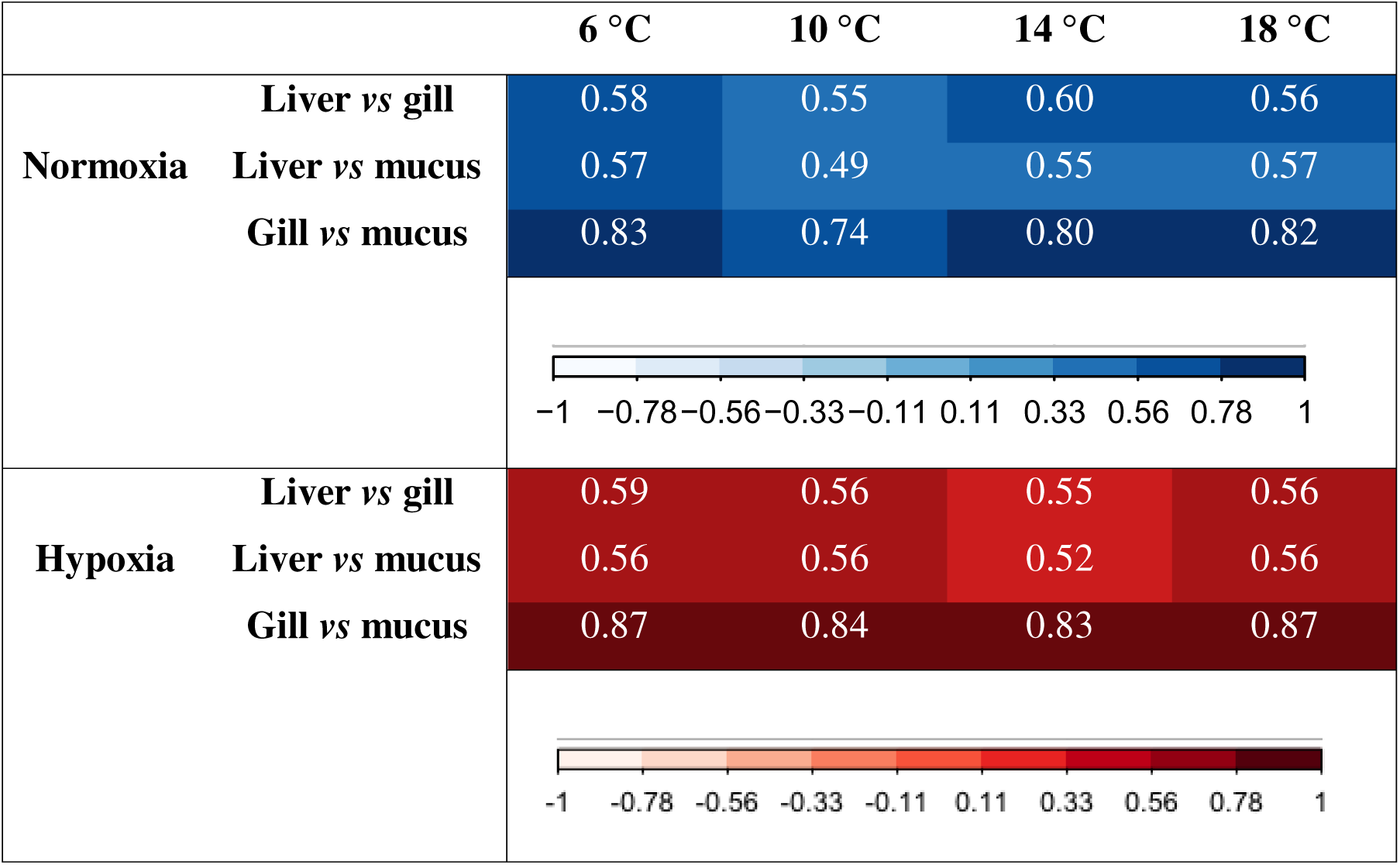
Pearson’s correlation coefficients of mRNA expression across liver, gill and mucus. Data presented are the compilation of shared genes between tissues with normalization to each ‘within-tissue’ endogenous control. Correlation plots were generated with ‘Corrplot’ v. 0.92 in R. Numbers and colours indicate the correlation coefficient.

## DISCUSSION

In the present study, the oxythermal transcriptional dependence of up to 46 genes across ten functional classifications was examined in three tissues of lake trout. Broadly, temperature exerted the greatest effect on the transcriptional profile with cellular stress markers further elevated under hypoxia, particularly at the highest tested temperature of 18 °C. Similarly, oxygen dependent thermal performance was noted at most tested temperatures with an overall reduction in CT_max_ in hypoxia treated fish compared to their normoxic counterparts. Following CT_max_, a drastic shift in the transcriptional profiles occurred, providing evidence of a shift toward a more extreme stress response (Jeffries *et al*., 2018). Together, these results highlight the use of sublethal transcriptional markers in assessing the physiological status and potential fitness consequences of wild cold-water fish populations that face exacerbated environmental perturbations with climate warming and deoxygenation.

Thermal acclimation improved thermal tolerance in lake trout, where with every 4 °C increase in acclimation temperature an ∼ 1 °C increase in CT_max_ occurred. The recorded temperatures are in agreeance with previous studies where CT_max_ was measured across similar acclimation ranges (8-19 °C) in four discrete populations of lake trout (Kelly *et al*., 2014). Further, thermal tolerance was oxygen dependent where the addition of hypoxia decreased CT_max_ in all tested temperatures, apart from 14 °C. This is suggestive of an energetic limitation in mounting a physiological response to acute warming, perhaps owing to both the reduced oxygen availability and increased tissue oxygen demand evoked by a CT_max_ trial. Both oxygen-dependent and oxygen-independent thermal tolerance phenotypes have been noted across fish species (Ern *et al*., 2016). Consequently, our finding of an oxygen-dependent phenotype at ecologically-relevant oxygen concentrations has important consequences in shaping lake trout distribution, particularly in the face of climate change.

Transcript abundance in the liver was strongly affected by acclimation temperature in lake trout. This is expected as the liver is a highly metabolically active tissue and the primary site of energy storage and mobilization (Volkoff & Rønnestad, 2020). Thus, when subjected to elevated temperatures and facing an increased metabolic demand (Kelly *et al*., 2014), the liver provides the energy required to sustain metabolic function (Volkoff & Rønnestad, 2020). Accordingly, the HSI decreased significantly with increasing acclimation temperature. An exacerbated decrease was noted in the hypoxia group at the most extreme temperatures (6, 18 °C), an indication of energy use. This temperature effect was noted for most measured transcripts despite the multitude of functional classifications, indicative of the role of temperature as a master regulator in ectotherms (Schulte *et al*., 2011). Across the six top contributors to the PCA, there was a consistent trend of elevated transcript abundance at 6 °C compared to all other temperatures in fish held in normoxia. With 10 °C considered within a preferred temperature range for lake trout (Christie & Regier, 1988), elevated transcripts may be indicative of energy mobilization as the fish aim to reestablish homeostasis at a cooler temperature. Similar alteration to hepatic metabolism was noted in bull trout (*S. confluentus*) acclimated to 6 °C (Best *et al*., 2024). Likewise, the decreased transcript abundance in elevated temperatures may be the result of energy reallocation where metabolism is downregulated, as has been reported for other salmonids (Beemelmanns *et al*., 2021; Jeffries *et al*., 2014). Lake trout exhibit a reduced metabolic scope when acclimated to 19 °C as compared to cooler acclimation temperatures (8-15°C; Kelly et al., 2014), suggesting that the optimal temperature has been surpassed and whole animal physiological processes are being negatively affected. The cellular mechanisms required to regain or maintain homeostasis are likely enacted prior to whole animal effects (Jeffries *et al*., 2018). Indeed, a significant downregulation of transcripts occurred here at 14 °C to a similar degree as at 18 °C, suggestive of a stress response already being mounted at 14 °C. A similar transcript profile was observed within the hypoxia group. However, transcript abundance at 6 and 10 °C was often similar perhaps suggesting that the fish in the hypoxia 6 °C group did not have the added energetic capacity to increase transcript abundance as did the normoxia group.

Despite an oxygen stressor in the range of the lower boundary for this species, oxygen dependent effects were not pronounced within liver transcripts. This corroborates a recent study in Atlantic salmon (*Salmo salar*) where the combined stressors of hypoxia and elevated temperature did not show additive effects on hepatic transcripts involved in the cellular stress response and instead clustered more readily by temperature treatment (Beemelmanns *et al*., 2021). Amongst the top six genes contributing to PC1 and PC3, three (*cs, casp3ab,* and *sod1*) can be considered markers of oxidative stress, the detoxification of which is another role enacted by the liver. *Cs* is the first step of the citric acid cycle, an integral aerobic biochemical process that yields energy from nutrients (Withers, 1992). As it is an aerobic process, oxygen limitation should limit *cs* activity, thereby increasing the need for increased activity or production of additional enzyme. While activity was not assessed within this dataset, transcript abundance suggests the latter is not occurring as oxygen dependent differences did not occur. *Casp3ab* is involved in the apoptosis cascade, which hypoxia can induce through inhibition of the electron transport chain (Ho *et al*., 2006), while *sod1* protects against damage caused by free radicals derived from oxygen that can be increased with hypoxic stress (Wang *et al*., 2018). No oxygen dependent changes were observed for *casp3ab*, while *sod1* showed a significant upregulation in normoxia only at the 6 °C acclimation, perhaps indicative of increased oxidative stress at this lower temperature. Similarly, no differences were observed between normoxia- and hypoxia-acclimated lake trout for other tested genes with roles in mediating hypoxic stress (e.g., *aldoaa, hk1, pgk1, scl2a1a*; SI Fig. 13). Together, this suggests that the acclimation period to each oxygen concentration was sufficient to reestablish homeostasis or the pre-existing levels of expressed proteins are sufficient to maintain physiological function under these reduced oxygen conditions.

The pronounced temperature dependence of transcript abundance observed in the liver was not evident in the gill of acclimated lake trout. Potentially, the energy mobilized by the liver supported maintained function of the gill, an organ responsible for multiple functions including osmo- and ionoregulation, nitrogenous waste excretion, and acid-base balance (Evans *et al*., 2005). At the interface between organism and environment, gills are often sensitive to environmental perturbation (Jeffries *et al*., 2021). Therefore, the relatively consistent expression profile across treatment groups suggests that the length of the acclimation period was sufficient to re-establish homeostasis or alternatively, these combinations were not stressful enough to alter gene expression. However, fish held at 18 °C and hypoxia did show a pronounced separation of gill transcripts in the PCA, as well as in gene expression profiles of aldolase (*aldoaa*), carnitine palmitoyltransferase 1(*cpt1a*)*, cs,* insulin-like growth factor 1 (*igf1*) and hydroxysteroid (20-beta) dehydrogenase 2 (*hsd20b2*). The first four of these genes are involved in metabolism, and the latter in the cellular stress response, demonstrating the metabolic demand evoked in this treatment group to maintain gill function. Interestingly, of the top six genes contributing to PC1 and PC2, three were consistent with those identified in the liver, *atp1b1, hspa4,* and *cam*. *Atp1b1* (i.e., sodium potassium ATPase [NKA]) is a ubiquitously expressed membrane transport protein responsible for establishing cellular electrochemical potential by exchanging three sodium ions outward for two potassium ions inward, making it important for osmoregulation (Evans *et al*., 2005). NKA also generates the ion gradient required for sodium-coupled transport in all tissues, where in the gill this may be linked to H^+^ or NH ^+^ (Evans *et al*., 2005). Given its crucial and ubiquitous cellular role, it is unsurprising that *atp1b1* contributed strongly to the profiles in both tissue types. Each of *hspa4* and *cam* also play ubiquitous roles as a heat shock protein chaperone and intracellular calcium binding protein, respectively, lending to their strong contribution to the PCA in both tissue types. Neither *hspa4* nor *cam* displayed distinct temperature or oxygen responses in the gill, although within the hypoxia group a significant upregulation of *hspa4* occurred at 18 °C, indicative of its role as a heat stress protectant (Kregel, 2002). Several other heat shock proteins (i.e., *hsp70a* and *hsp7c*) were also upregulated with increasing temperature, much like has been shown in other teleosts (Huang *et al*., 2022; Jeffries *et al*., 2014), as evidence of continued heat stress over the duration of the experiment.

Fewer genes and individuals contributed to the PCA for the mucus samples, leading to the absence of clear separation based upon treatment. Instead, a general trend of decreased transcript abundance was observed at 10 °C compared to all other temperatures, with marked exacerbation within the hypoxia treatment at 10 °C. The elevation in transcripts may be indicative of a cellular stress response to non-optimal temperatures; either cool or warm. Akin to the liver and gill, *cam* and *atp1b1* were two of the top contributing genes to the overall expression profiles in mucus, hinting at their ubiquitous importance across tissues. Similarly, *cirpba*, a gene involved in RNA stabilization and translation efficiency (Jeffries *et al*., 2014), was a top contributor in the gill and the mucus, with stronger temperature-dependent expression observed in the mucus. Both *cam* and *gstp1* had significant oxygen dependent differences at 10°C. Fittingly, each of these genes can be activated in response to oxidative stress and play roles in mitigating redox stress (Franklin *et al*., 2006; Russell & Richardson, 2023).

Acute heat stress to lake trout upper thermal limits elicited a marked response in both liver and gill, particularly in the 18 °C-hypoxia acclimated fish. As in control-acclimated fish, this response did not extend to the mucus samples. Despite this, many common genes contributed to the top six responders in each tissue, with some akin to those found in the acclimation trials (e.g., *cirpba, sod1, gstp1*). Notably, markers for lipid metabolism were significantly increased at 18 °C-hypoxia, including hepatic *cpt1a* and branchial lipoprotein lipase (*lpl*) and fatty acid synthase (*fasn*). Other glycolytic enzymes were significantly affected including *pgk* and *aldoaa*, again suggestive of increased metabolic flux under this most stressful condition. Similar increases in lipid metabolism were found in liver and muscle samples of bull trout acclimated to high temperature (21 °C; Best et al., 2024), marking the mobilization of energetic stores to fuel homeostatic maintenance in the face of environmental stress. These underlying transcriptional differences may contribute to the decline in temperature tolerance under hypoxic conditions as assessed by CT_max_.

Finally, a distinct shift in mucus transcript abundance in acclimation temperatures outside the preferred 10 °C highlights the potential use of mucus swabs as a non-lethal tool for management and conservation of fish. The method used herein of collecting mucus using swabs likely has RNA contributions from multiple sources including skin, scales, and potentially blood. However, the collective ‘mucus’ sample yields transcriptional responses that correlate well with those of the gill as verified by a Pearson’s correlation coefficient of *r* = 0.74-0.87 compared to liver *vs* mucus (*r* = 0.49-0.57) or liver *vs* gill correlations (*r* = 0.55-0.60). This is unsurprising given their similar contact with the external milieu where they are both metabolically active tissues acting as external barriers (innate immunity) and participating in respiration and excretion (Evans 2004; Raposo De Magalhães et al., 2023). Previous studies have demonstrated that both biotic and abiotic stressors can alter mucus production and composition in teleosts (e.g., (Andrzejczyk *et al*., 2022; Easy & Ross, 2010; Reverter *et al*., 2018; Terova *et al*., 2011; Zheng *et al*., 2021). Indeed, within our own data we note significant differences in the transcriptional profile of genes involved in the protection of the proteome from stress (i.e., *hsp70a* and *hsp7c*; SI Fig. 18) depending upon acclimation treatment. Further, the strength of correlation increases outside of the preferred oxythermal habitat at 10 °C-normoxia, indicative of similarly altered transcriptional profiles in response to environmental stressors. Together, this supports the use of transcriptional profiles as indicators of stress in wild fish (Jeffries et al. 2018(Andrzejczyk *et al*., 2022), with mucus being an excellent tissue for conducting non-lethal sampling of culturally and ecologically important fish species.

## ACKNOWLEDGEMENTS

The authors would like to acknowledge the aid of Carol Best and Hossein Haghighi in learning how to operate the QuantStudio 12K Flex System, and Linh Miller, Carolyn Vandervelde, Stefanie Kornberger, and Holly Simpson for assistance with sampling and the CT_max_ estimates. Additional thanks to Kerry Wautier for help with fish holding and husbandry and Ian Bouyoucos for helpful discussions and proof-reading of an earlier draft of this manuscript.

## AUTHOR CONTRIBUTIONS

AMW – Methodology, Validation, Formal Analysis, Investigation, Writing-Original Draft, Writing – Review & Editing, Visualization

ALC - Methodology, Validation, Formal Analysis, Investigation, Writing – Review & Editing, Visualization

TCD – Conceptualization, Methodology, Validation, Investigation, Writing – Review & Editing

EE – Conceptualization, Resources, Supervision, Methodology, Validation, Investigation, Writign – Review & Editing, Funding Acquisition

KMJ – Conceptualization, Resources, Supervision, Project Administration, Funding Acquisition, Writing – Review & Editing

## DATA AVAILABILITY

Data are available in the supplemental information.

