## Supporting Information for "Cumulative effects of high temperature and low dissolved oxygen alter the acute thermal tolerance and cellular stress response in lake trout"

A comparison between the time-separated controls (i.e., normoxia and normoxia_2) was first conducted for each type of analysis to determine whether temporal differences occurred between the normoxia and hypoxia experimental trials. Regarding $CT_{max}$ data, no significant differences were observed between the time-separated controls as determined by a Wilcoxon rank sum test ($p = 0.25$), so the latter was dropped from the overall comparison. Similarly, comparisons of gene transcript abundance were compared between the two time-separated controls. In the liver, only *cry1ab* showed differential expression indicating that 97.96% of investigated genes were unaffected across time within the same treatment (SI Fig. 1). In the gill, no genes were found to be differentially expressed (SI Fig. 2), while in the mucus *hsp70a* showed time-dependent expression (SI Fig. 3), indicating that 100% and 96.88% of genes remain unchanged over time, respectively. Within the $CT_{max}$ fish, a similar comparison was made where liver showed differential expression between time-separated controls for only three genes (*cpt1a, ghr, lipeab*; SI Fig. 4), while gill (SI Fig. 5) and mucus (SI Fig. 6) showed no differentially expressed genes between these controls. Given the relative stability of most genes across these time-separated controls, the normoxia group that was collected alongside the hypoxic trials (i.e., normoxia_2) was omitted from all further analysis.

Transcript abundance was first analysed using a naïve model of the MCMC.qpcr package to establish which housekeeping genes remained stable across treatments to be used as priors in the informed models. This differed across tissues with *rpl7* used for the liver (SI Fig. 7), *rpl7, rpl13a,* and *rps9* used in the gill (SI Fig. 8), and *rpl7, rpl13a, rps9*, and *ef1α* used in the mucus (SI Fig. 9) informed models generated for the control, acclimated fish. For data from the $CT_{max}$ fish, *rpl7* was the sole housekeeping gene used for the liver (SI Fig. 10) and gill (SI Fig. 11), while all four genes were used to inform the mucus models (SI Fig. 12).

**SI Table 1. Genes on the salmonid stress-response transcriptional profiling chip and their associated functional classification.**

| Gene Name | Functional Classification | Liver Efficiency | Gill Efficiency | Mucus Efficiency |
|---|---|---|---|---|
| Caspase3AB (*casp3AB*) | Apoptosis | 1.894 | 1.908 | 1.925 |
| Caspase9 (*casp9*) | Apoptosis | 1.766 | 1.891 | 1.827 |
| Charged multivesicular body 5 (*chmp5AB*) | Apoptosis | 1.618 | NA | 1.622 |
| Programmed cell death protein 10 (*pdcd10*) | Apoptosis | 1.981 | 2.004 | 2.017 |
| CLOCK1A (*clock1a*) | Circadian rhythm | 1.933 | 1.903 | 1.851 |
| Cryptochrome circadian regulator 1 (*cry1ab*) | Circadian rhythm | 1.93 | 1.889 | NA |
| Catalase (*cat*) | Detoxification | 1.866 | 1.842 | NA |
| Glutathione S-transferase pi 1 (*gstp1*) | Detoxification | 1.969 | 2.024 | 1.902 |
| Nuclear factor erythroid 2-related factor 2 (*nfe2l2a*) | Detoxification | 2.055 | 2.062 | 2.034 |
| Superoxide dismutase 1 (*sod1*) | Detoxification | 2.137 | 2.045 | 1.991 |
| Superoxide dismutase 2 (*sod2*) | Detoxification | 2.097 | 2.119 | 2.037 |
| Aldolase (*aldoaa*) | Growth/ metabolism | 2.023 | 2.192 | 2.014 |
| Carnitine palmitoyl transferase 1 (*cpt1a*) | Growth/ metabolism | 2.113 | 2.085 | NA |
| Citrate synthase (*cs*) | Growth/ metabolism | 1.994 | 2.053 | 1.99 |
| Cathepsin D (*ctsd*) | Growth/ metabolism | 1.971 | 1.975 | 1.926 |
| Fatty acid synthase (*fasn*) | Growth/ metabolism | 2.00 | 1.893 | 1.93 |
| Growth hormone receptor (*ghr*) | Growth/ metabolism | 1.886 | 1.873 | NA |

| | | | | |
|---|---|---|---|---|
| Insulin-like growth factor 1 (*igf1*) | Growth/ metabolism | 1.863 | 1.897 | NA |
| Insulin-like growth factor 2 (*igf2*) | Growth/ metabolism | 1.871 | 1.951 | NA |
| Insulin-like growth factor binding protein 1 (*igfbp1*) | Growth/ metabolism | 1.696 | NA | NA |
| Lactate dehydrogenase A (*ldha*) | Growth/ metabolism | NA | 2.049 | NA |
| Lactate dehydrogenase B (*ldhb*) | Growth/ metabolism | 1.985 | 2.11 | 1.999 |
| Lipase (*lipeab*) | Growth/ metabolism | 2.134 | 2.11 | 2.085 |
| Lipoprotein lipase (*lpl*) | Growth/ metabolism | 2.064 | 2.118 | NA |
| Phosphoenolpyruvate carboxykinase 1 (*pck1*) | Growth/ metabolism | NA | NA | NA |
| Phosphoglycerate kinase (*pgk*) | Growth/ metabolism | 1.928 | 1.984 | 1.95 |
| 5’ AMP-activated protein kinase (*ampka1*) | Hypoxia/ metabolism | 2.01 | 2.035 | 2.042 |
| Hypoxia inducible factor 1 alpha (*hif1a*) | Hypoxia/ metabolism | NA | NA | NA |
| Hexokinase 1 (*hk1*) | Hypoxia/ metabolism | 1.871 | 1.941 | 1.93 |
| Leptin receptor (*lepr*) | Hypoxia/ metabolism | NA | 1.979 | NA |
| Glucose transporter (*slc2a1a*) | Hypoxia/ metabolism | 1.618 | 1.748 | 1.757 |
| Vascular endothelial growth factor (*vegf*) | Hypoxia/ metabolism | NA | 1.884 | NA |
| Calmodulin (*cam*) | Immune | 2.056 | 2.241 | 2.025 |
| Major histocompatibility complex I (*mhci*) | Immune | 2.154 | 2.25 | 2.1 |

| | | | | |
|---|---|---|---|---|
| Signal transducer and activator of transcription 1 (*stat1*) | Immune | 1.925 | 1.987 | 1.781 |
| Sodium potassium ATPase beta subunit (*atp1b1*) | Osmoregulation | 1.879 | 1.954 | 1.865 |
| Cold inducible RNA binding protein a (*cirbpa*) | Stress response | 2.186 | 2.196 | 1.997 |
| Glucocorticoid receptor 2 (*gr2*) | Stress response | 1.945 | 1.915 | 1.926 |
| Hydroxysteroid 11-beta dehydrogenase 2 (*hsd11b2*) | Stress response | 1.941 | 2.044 | NA |
| Hydroxysteroid 20-beta dehydrogenase 2 (*hsd20b2*) | Stress response | 1.951 | 2.015 | NA |
| Heat shock transcription factor 1 (*hsf1*) | Stress response | 2.108 | 2.097 | NA |
| Heat shock 70a (*hsp70a*) | Stress response | 1.944 | 1.953 | 1.946 |
| Heat shock ccognate 71 kDa protein (*hsp7c*) | Stress response | 2.029 | 1.99 | 2.0 |
| Heat shock protein 90 inducible (*hsp90aa*) | Stress response | NA | NA | NA |
| Heat shock protein 90 constitutive (*hsp90ba*) | Stress response | 1.715 | 1.75 | 1.705 |
| Heat shock protein family a member 4 (*hspa4*) | Stress response | 2.048 | 2.041 | 1.981 |
| Matrix metallopeptidase 2 constitutive (*mmp2*) | Stress response | 2.065 | 2.15 | NA |
| Matrix metallopeptidase 9 inducible (*mmp9*) | Stress response | 1.905 | 2.058 | 1.94 |
| Mineralocorticoid receptor (*mr*) | Stress response | 1.907 | 1.962 | 1.877 |

| | | | | |
|---|---|---|---|---|
| Metallothionein A (*mtA*) | Stress response | 2.025 | 1.878 | 2.044 |
| Metallothionein B (*mtB*) | Stress response | NA | NA | NA |
| Serpin family H1 (*serpinh1*) | Stress response | 1.92 | 1.957 | NA |
| Elongation factor 1 alpha (*ef1a*) | Endogenous control | 2.008 | 2.037 | 1.96 |
| Ribosomal protein 13A (*rpl13a*) | Endogenous control | 2.087 | 2.123 | 1.971 |
| Ribosomal protein 7 (*rpl7*) | Endogenous control | 1.894 | 1.961 | 1.856 |
| Ribosomal protein s9 (*rps9*) | Endogenous control | 1.982 | 2.027 | 1.954 |

**SI Table 2. Temperature and hypoxia treatment effects on hepatosomatic index (HSI) of acclimated and/or acutely heat stressed ($CT_{max}$) lake trout (*Salvelinus namaycush*).** Different letters denote significant pairswise differences as estimated using the *R* package *emmeans* v 1.9.0 with lowercase letters comparing control acclimated fish and uppercase letters comparing $CT_{max}$ fish.

| **Acclimation group** | **HSI** (mean ± s.e.m.) | **Estimated Marginal Mean ± s.e.m.** | **Model Output** | | | |
|---|---|---|---|---|---|---|
| Normoxia (N), 6 °C | 1.08 ± 0.06[a] | 1.08 ± 0.052 | | DF | F value | *p* |
| N 10 °C | 0.88 ± 0.04[a] | 0.88 ± 0.052 | | | | |
| N 14 °C | 0.96 ± 0.06[a] | 0.96 ± 0.052 | Oxygen | 1 | 17.51 | < 0.001 |
| N 18 °C | 0.99 ± 0.09[a] | 0.99 ± 0.058 | Temperature | 3 | 6.40 | < 0.001 |
| Hypoxia (H), 6 °C | 0.89 ± 0.03[a] | 0.89 ± 0.052 | Oxygen:Temperature | 3 | 6.95 | < 0.001 |
| H 10 °C | 0.91 ± 0.05[a] | 0.91 ± 0.052 | Residuals | 108 | | |
| H 14 °C | 0.92 ± 0.04[a] | 0.92 ± 0.054 | | | | |
| H 18 °C | 0.57 ± 0.03[b] | 0.56 ± 0.052 | | | | |
| N_2 (10 °C) | 0.91 ± 0.04 | | Wilcoxon (Normoxia, 10°C vs Normoxia_2) | | NA | $p = 0.26$ |
| $CT_{max}$ N 6 °C | 1.17 ± 0.06 | 1.18 ± 0.048[A] | | DF | F value | *p* |

| | | | | | | |
|---|---|---|---|---|---|---|
| $CT_{max}$ N 10 °C | 0.88 ± 0.04 | 0.89 ± 0.048[B] | Oxygen | 1 | 12.04 | < 0.001 |
| $CT_{max}$ N 14 °C | 0.92 ± 0.06 | 0.92 ± 0.048[B] | Temperature | 3 | 4.00 | < 0.01 |
| $CT_{max}$ N 18 °C | 1.01 ± 0.06 | 1.01 ± 0.048[B] | Oxygen:Temperature | 3 | 3.47 | 0.02 |
| $CT_{max}$ H 6 °C | 0.90 ± 0.04 | 0.90 ± 0.048[B] | Residuals | 110 | | |
| $CT_{max}$ H 10 °C | 0.90 ± 0.04 | 0.91 ± 0.048[B] | | | | |
| $CT_{max}$ H 14 °C | 0.87 ± 0.04 | 0.87 ± 0.048[B] | | | | |
| $CT_{max}$ H 18 °C | 0.83 ± 0.04 | 0.83 ± 0.052[B] | | | | |

SI: Cumulative effects of high temperature and low dissolved oxygen alter the acute thermal tolerance and cellular stress response in lake trout

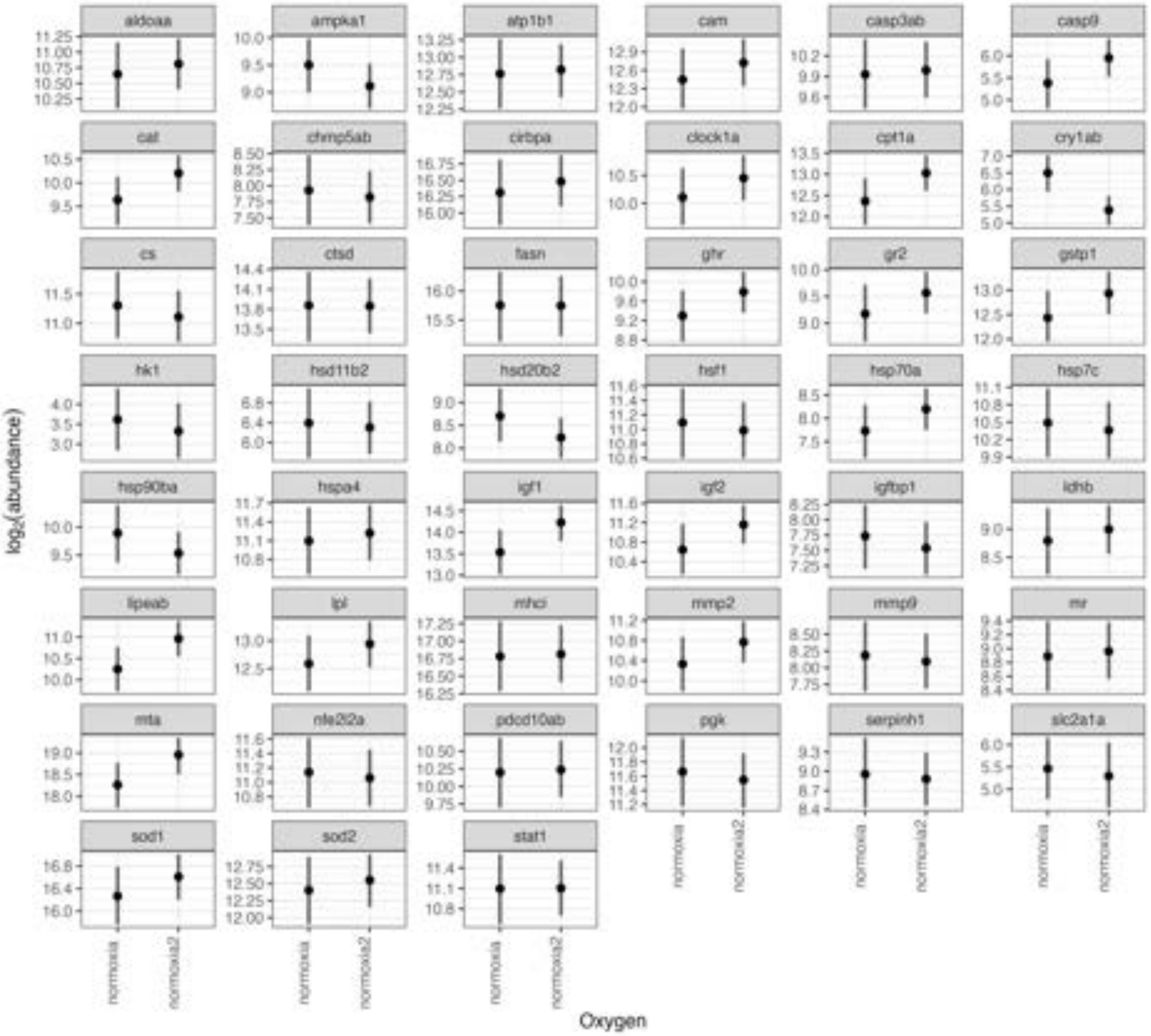


**SI Figure 1. Comparison of the normoxia 10 °C liver mRNA transcript abundance at the two experimental trial times in control, acclimated lake trout (*Salvelinus namaycush*).** The same treatment was applied at two different acclimation times (normoxia and normoxia_2, the latter conducted during the hypoxia trials) to ensure consistency across time in expression. All genes, except for cry1ab, were not differentially expressed between times ($n$ = 6-15).

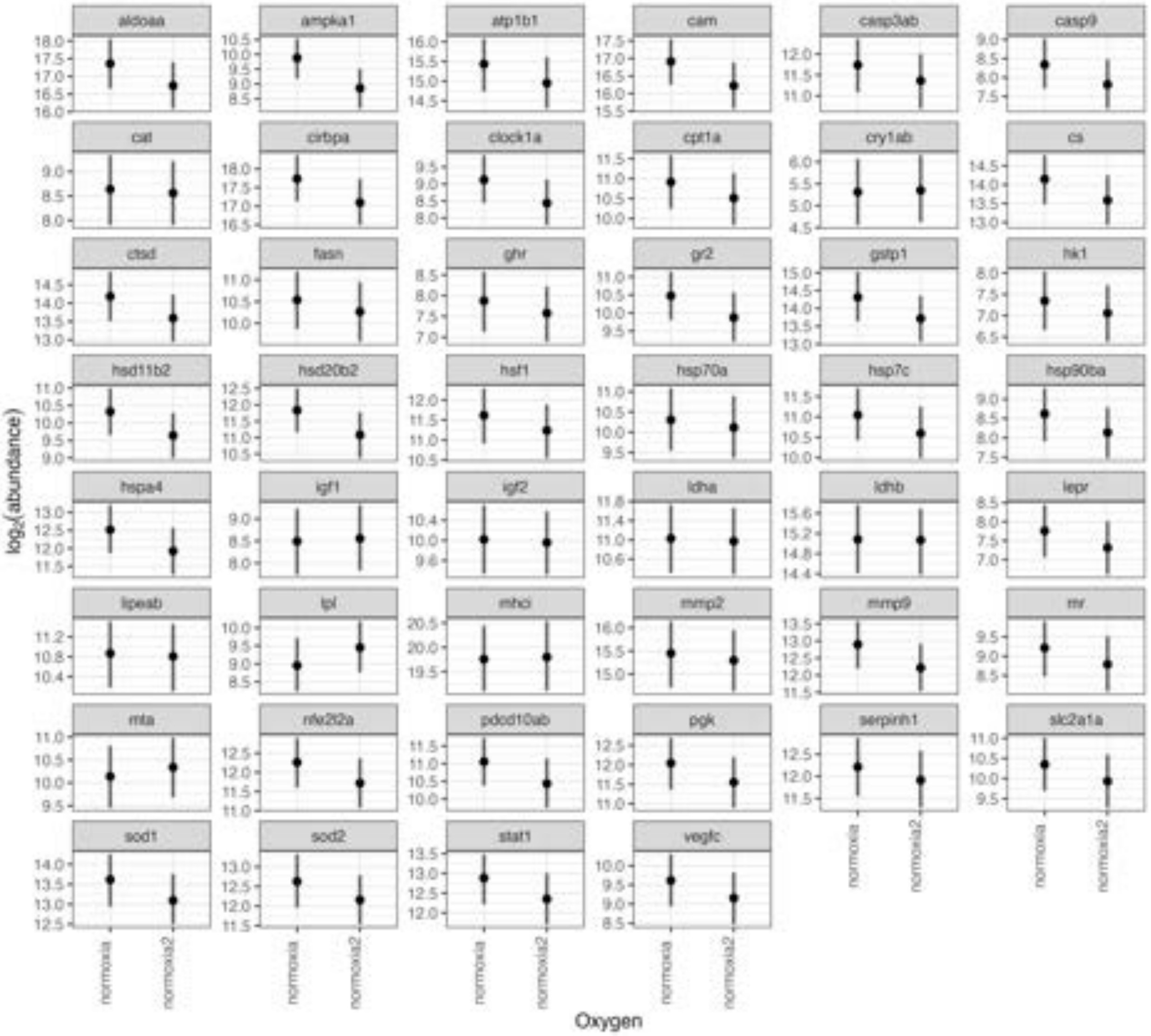


**SI Figure 2. Comparison of the normoxia 10 °C gill mRNA transcript abundance between the two experimental trial times in control, acclimated lake trout (*Salvelinus namaycush*).** The same treatment was applied at two different acclimation times (normoxia and normoxia_2, the latter conducted during the hypoxia trials) to ensure consistency across time in expression. No differences were observed between the two groups in the gill ($n$ = 8-13).

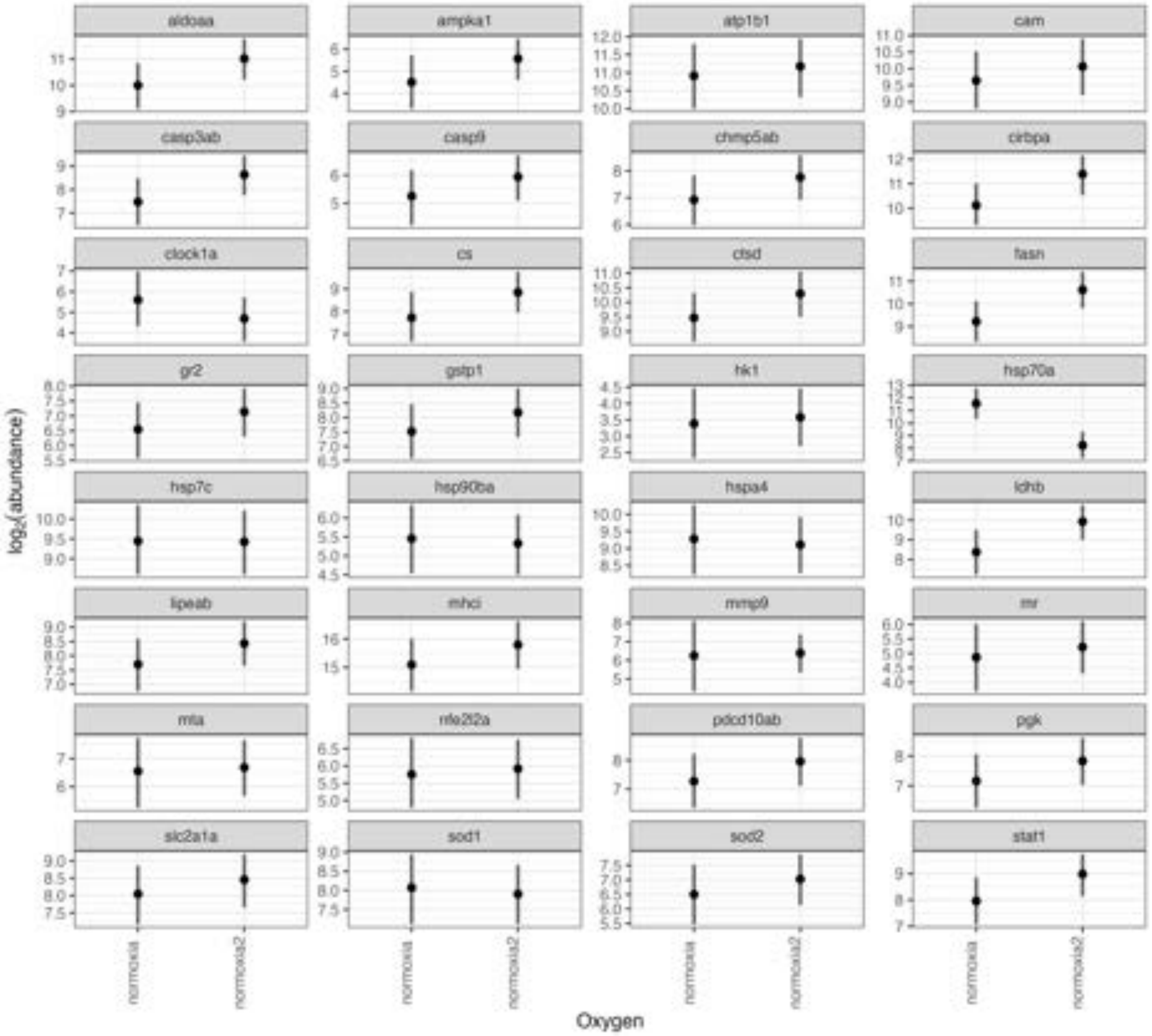


**SI Figure 3. Comparison of the normoxia 10 °C mucus mRNA transcript abundance at two different experimental trial times in control, acclimated lake trout (*Salvelinus namaycush*).** The same treatment was applied at two different acclimation times (normoxia and normoxia_2, the latter conducted during the hypoxia trials) to ensure consistency across time in expression. No differences were observed except for *hsp70a* ($n$ = 4-9).

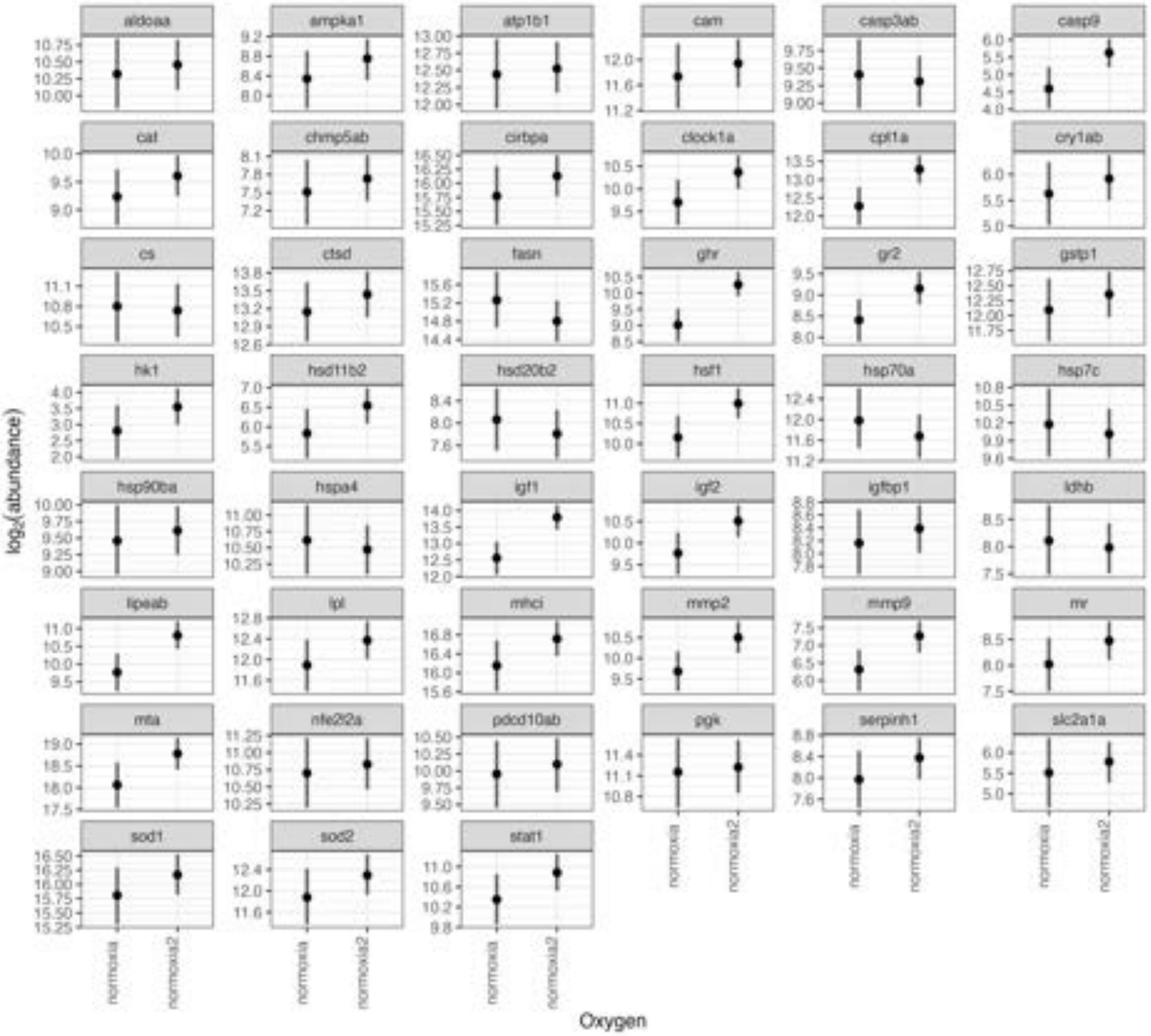


**SI Figure 4. Comparison of the normoxia 10°C liver mRNA transcript abundance at the two experimental trial times in lake trout (*Salvelinus namaycush*) after $CT_{max}$ trials.** The same treatment was applied at two different acclimation times (normoxia and normoxia_2, the latter conducted during the hypoxia trials) to ensure consistency across time in expression. All genes, except for *cpt1a*, *ghr*, and *lipeab* were not differentially expressed between times (*n* = 6-12).

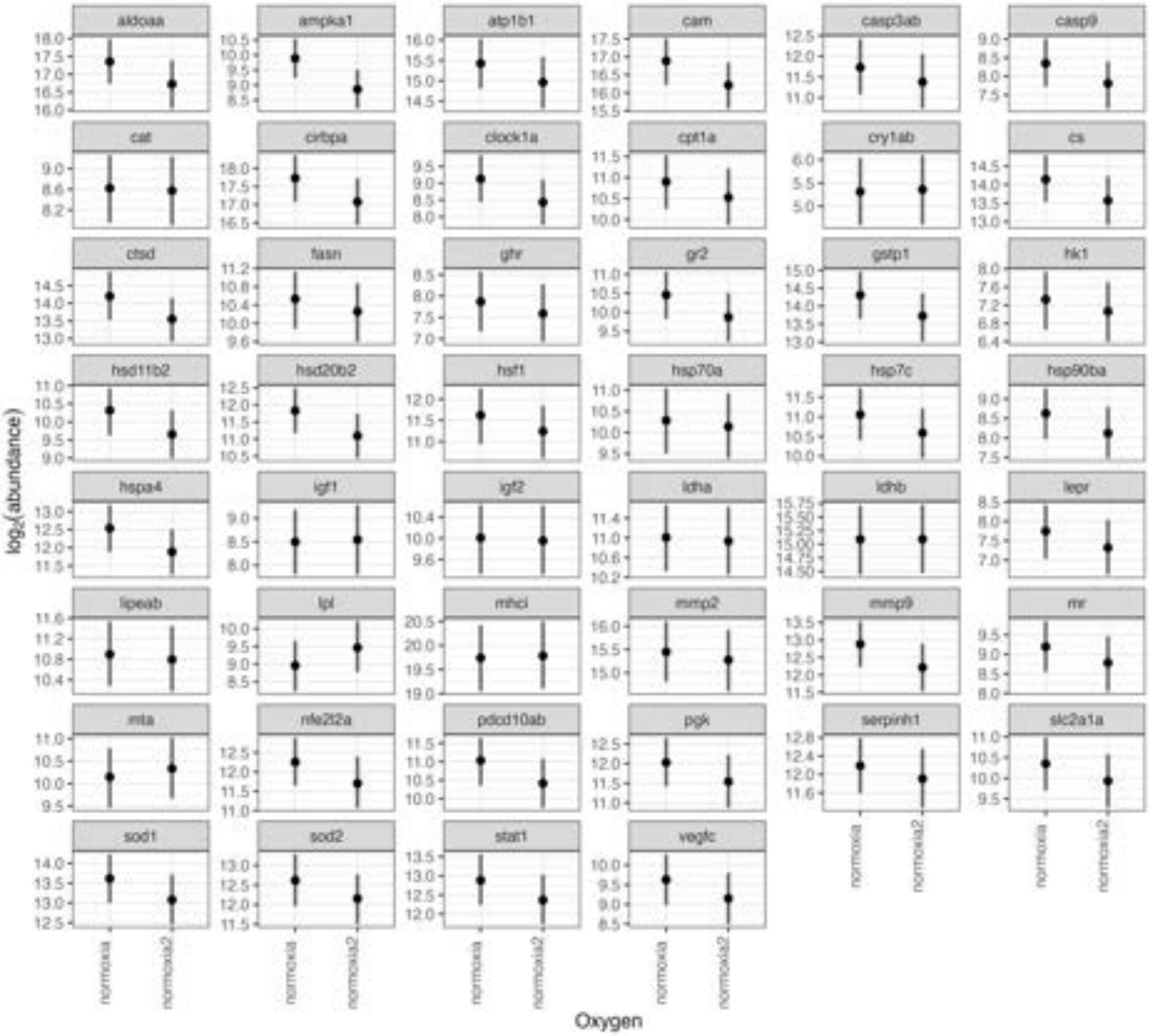


**SI Figure 5. Comparison of the normoxia 10 °C gill mRNA transcript abundance at the two experimental trial times in lake trout (*Salvelinus namaycush*) after $CT_{max}$ trials.** The same treatment was applied at two different acclimation times (normoxia and normoxia_2, the latter conducted during the hypoxia trials) to ensure consistency across time in expression. No genes were found to be differentially expressed (*n* = 8-11).

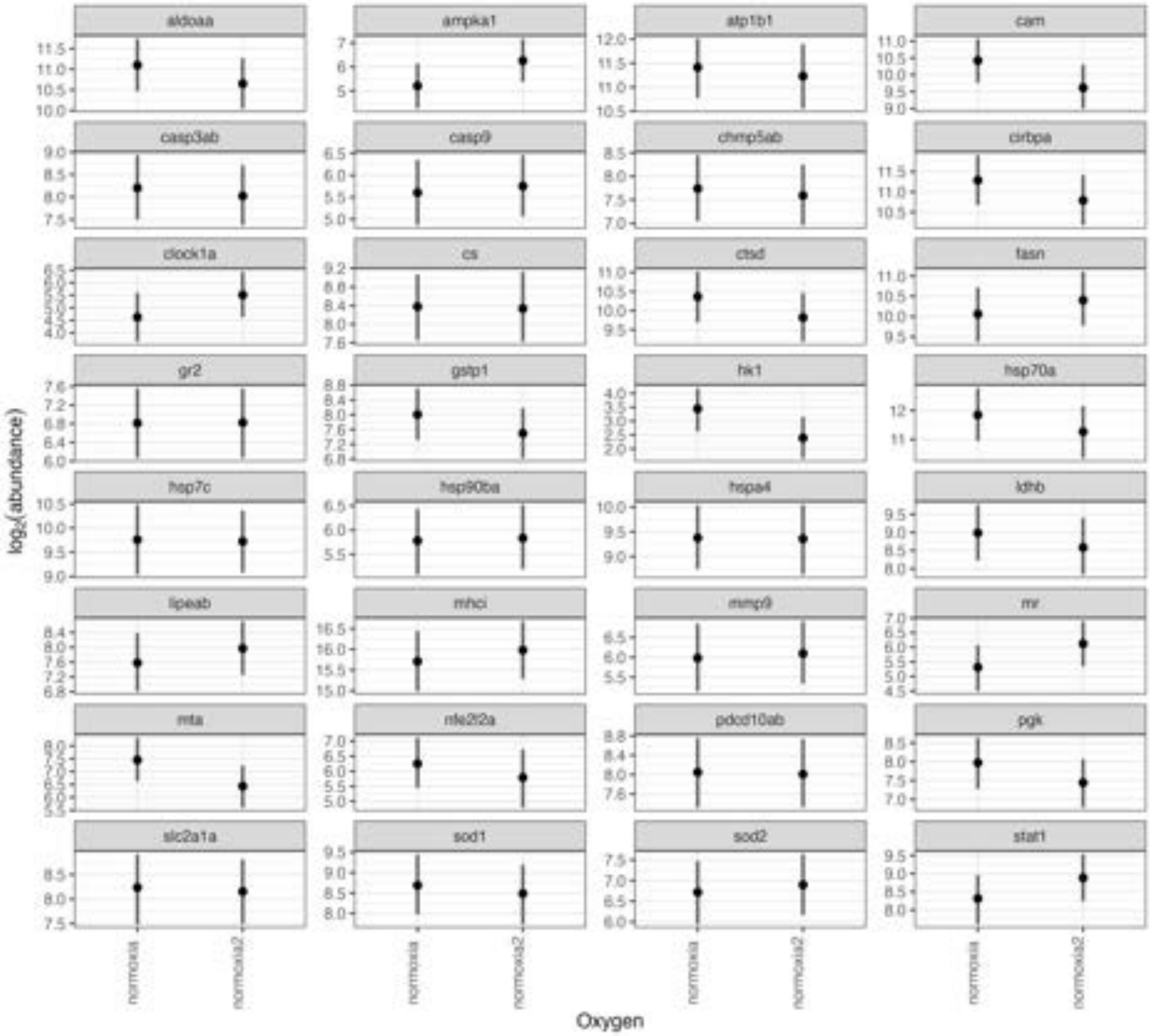


**SI Figure 6. Comparison of the normoxia 10 °C mucus mRNA transcript abundance at the two experimental trial times in lake trout (*Salvelinus namaycush*) after $CT_{max}$ trials.** The same treatment was applied at two different acclimation times (normoxia and normoxia_2, the latter conducted during the hypoxia trials) to ensure consistency across time in expression. No tested genes were found to be differentially expressed between the two groups ($n$ = 5-10).

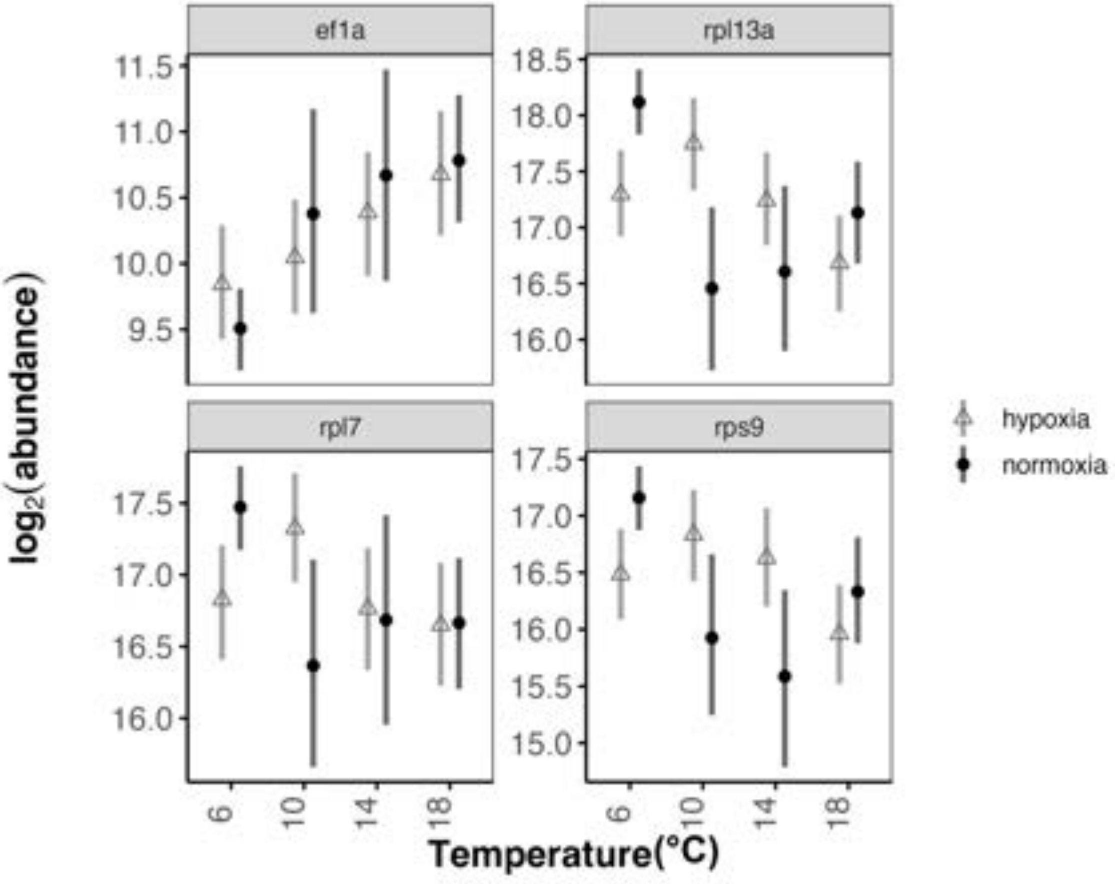


**SI Figure 7. Naïve MCMC.qpcr model of hepatic transcript abundance housekeeping genes across temperature acclimations and oxygen concentrations in control, acclimated lake trout (*Salvelinus namaycush*).** *Rpl7* was the only housekeeping gene used as a prior in the informed models contributing to Figure 2. Values from lake trout ($n$ = 6-15) acclimated to four temperatures (6, 10, 14, 18 °C) and two oxygen concentrations (normoxia: 9.93 ± 1.23 mg $L^{-1}$; hypoxia: 6.18 ± 0.44 mg $L^{-1}$).

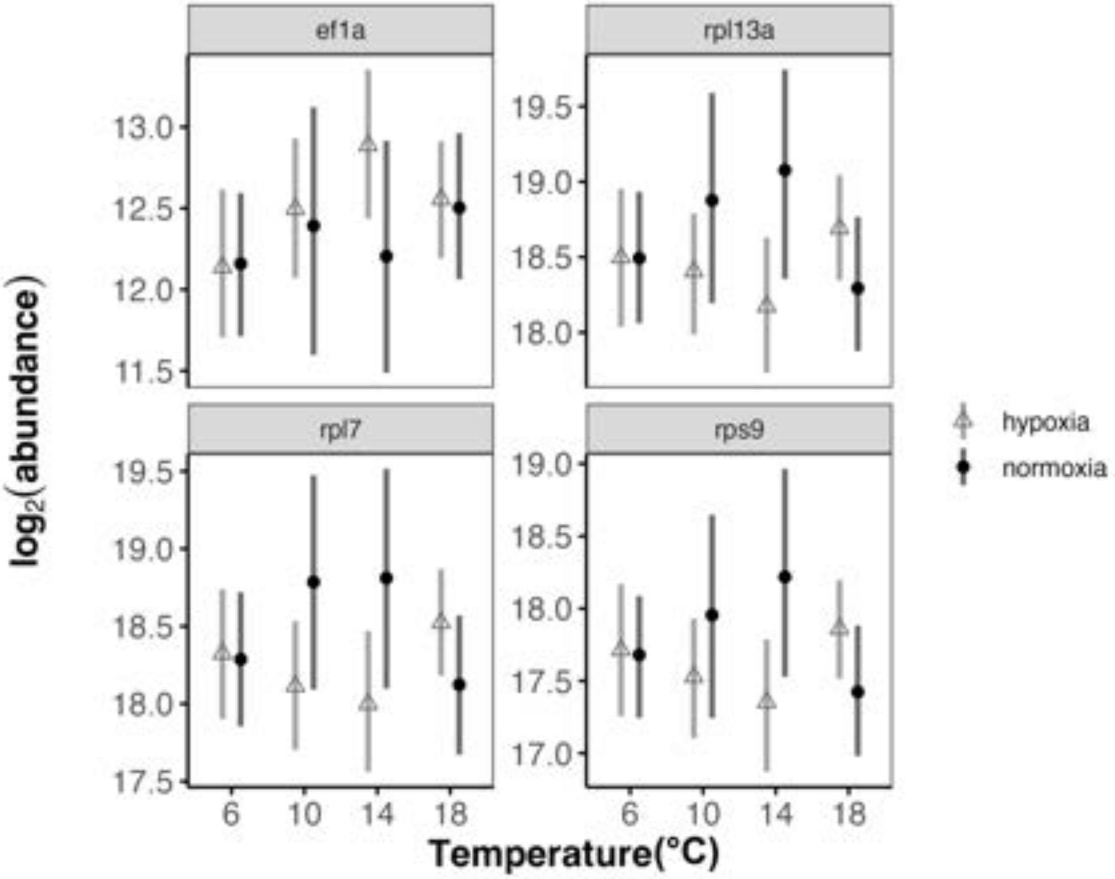


**SI Figure 8. Naïve MCMC.qpcr model of branchial transcript abundance of housekeeping genes across temperature acclimations and oxygen concentrations in control, acclimated lake trout (*Salvelinus namaycush*).** *Rpl13a*, *rpl7*, and *rps9* were used has priors in the informed model that contributed to Figure 3. Values from lake trout (*n* = 8-13) acclimated to four temperatures (6, 10, 14, 18 °C) and two oxygen concentrations (normoxia: 9.93 ± 1.23 mg $L^{-1}$; hypoxia: 6.18 ± 0.44 mg $L^{-1}$).

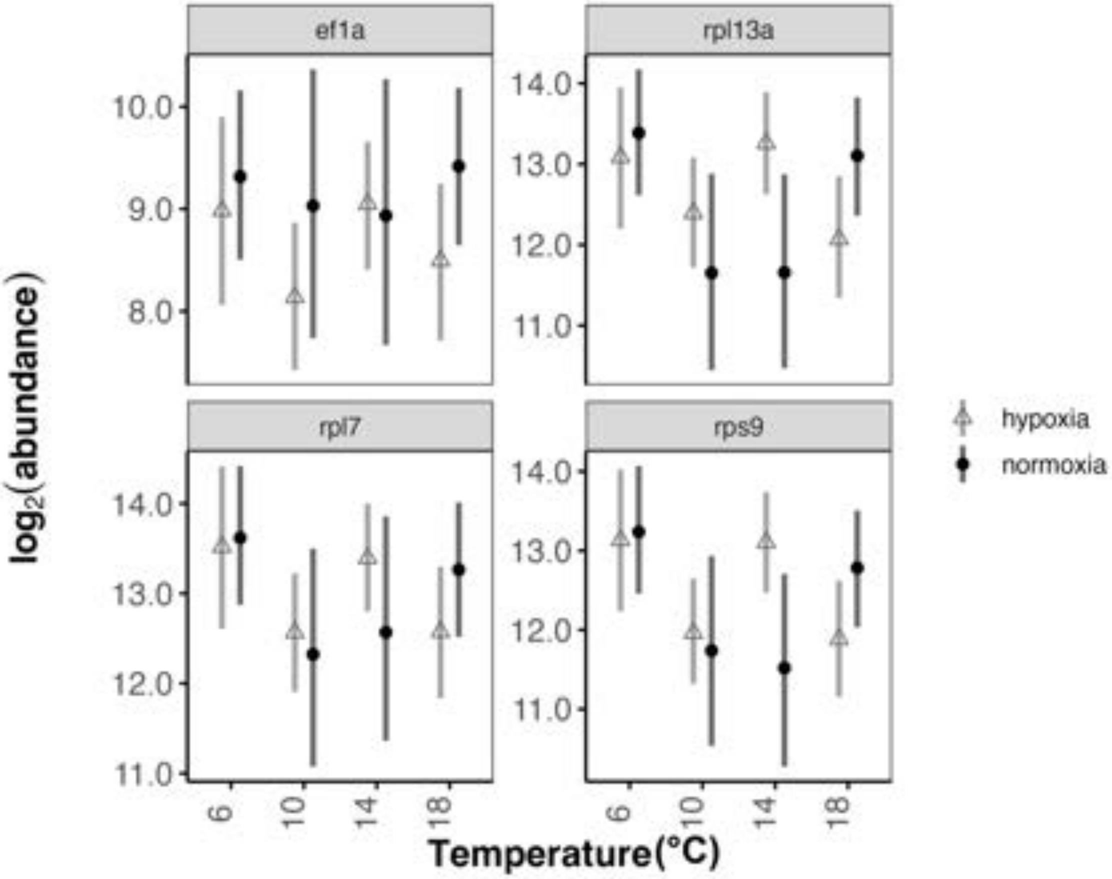


**SI Figure 9. Naïve MCMC.qpcr model of transcript abundance of housekeeping genes in the mucus across temperature acclimations and oxygen concentrations in control, acclimated lake trout (*Salvelinus namaycush*).** All four housekeeping genes were used as priors in the informed model used to generate Fig. 4. Values from lake trout ($n$ = 4-9) acclimated to four temperatures (6, 10, 14, 18 °C) and two oxygen concentrations (normoxia: 9.93 ± 1.23 mg $L^{-1}$; hypoxia: 6.18 ± 0.44 mg $L^{-1}$).

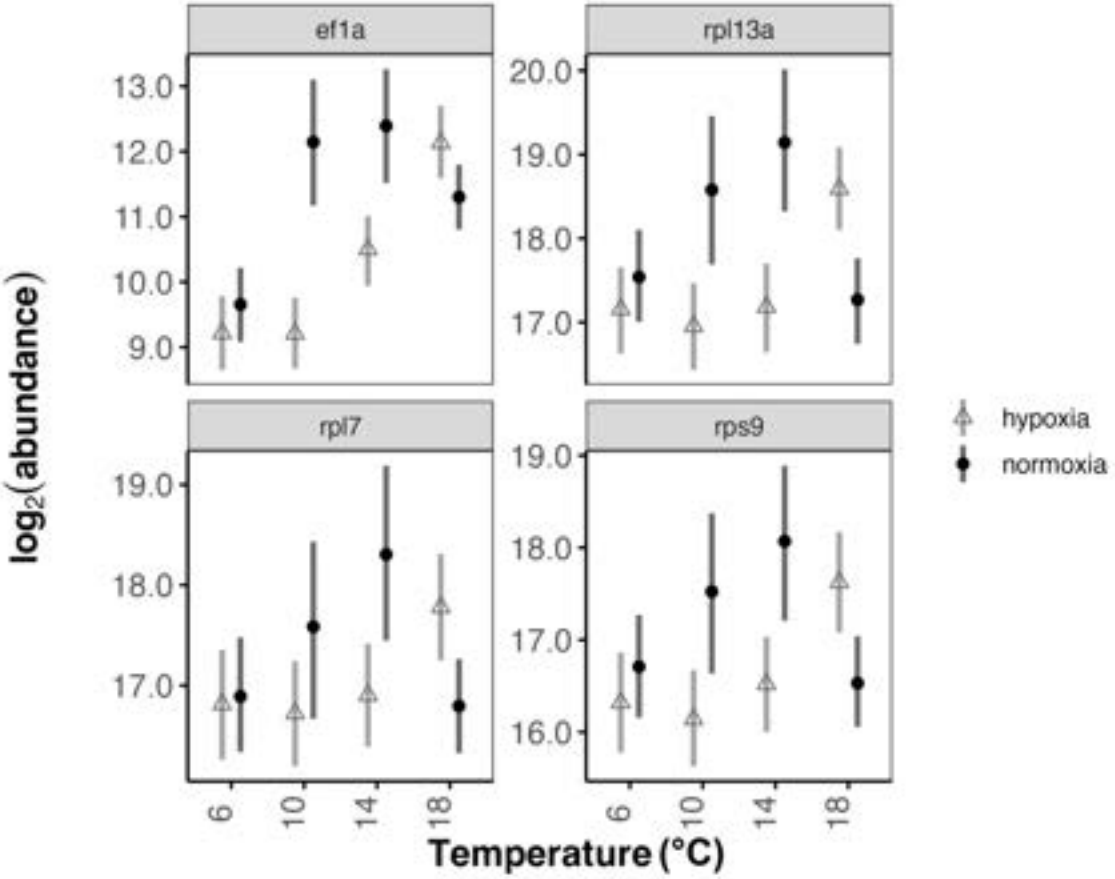


**SI Figure 10. Naïve MCMC.qpcr model of transcript abundance of housekeeping genes in the liver across temperature acclimations and oxygen concentrations in $CT_{max}$ lake trout (*Salvelinus namaycush*).** Only *rpl7* was used in the informed model. Values from lake trout ($n$ = 6-12) acclimated to four temperatures (6, 10, 14, 18°C) and two oxygen concentrations (normoxia: 9.93 ± 1.23 mg $L^{-1}$; hypoxia: 6.18 ± 0.44 mg $L^{-1}$).

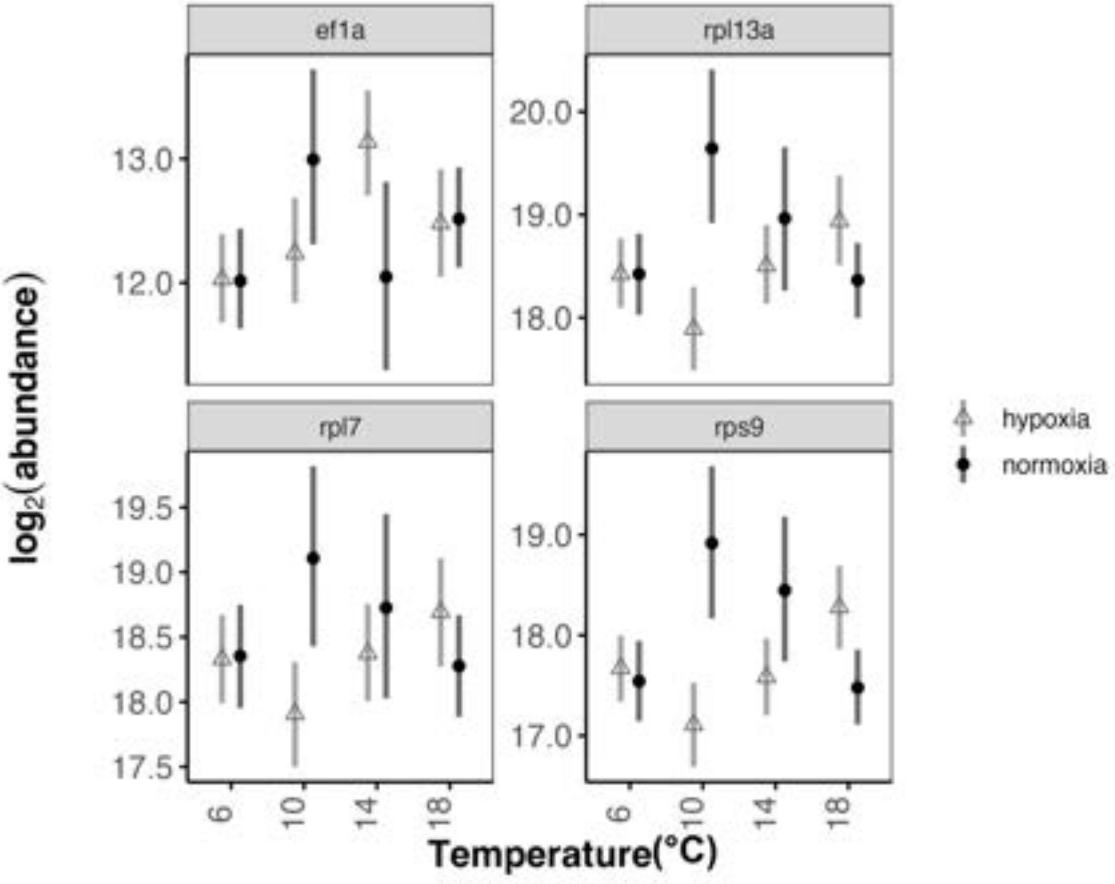


**SI Figure 11. Naïve MCMC.qpcr model of transcript abundance of housekeeping genes in the gill across temperature acclimations and oxygen concentrations in $CT_{max}$ lake trout (*Salvelinus namaycush*).** Only *rpl7* was used as a prior in the informed model. Values from lake trout ($n$ = 8-11) acclimated to four temperatures (6, 10, 14, 18°C) and two oxygen concentrations (normoxia: 9.93 ± 1.23 mg $L^{-1}$; hypoxia: 6.18 ± 0.44 mg $L^{-1}$).

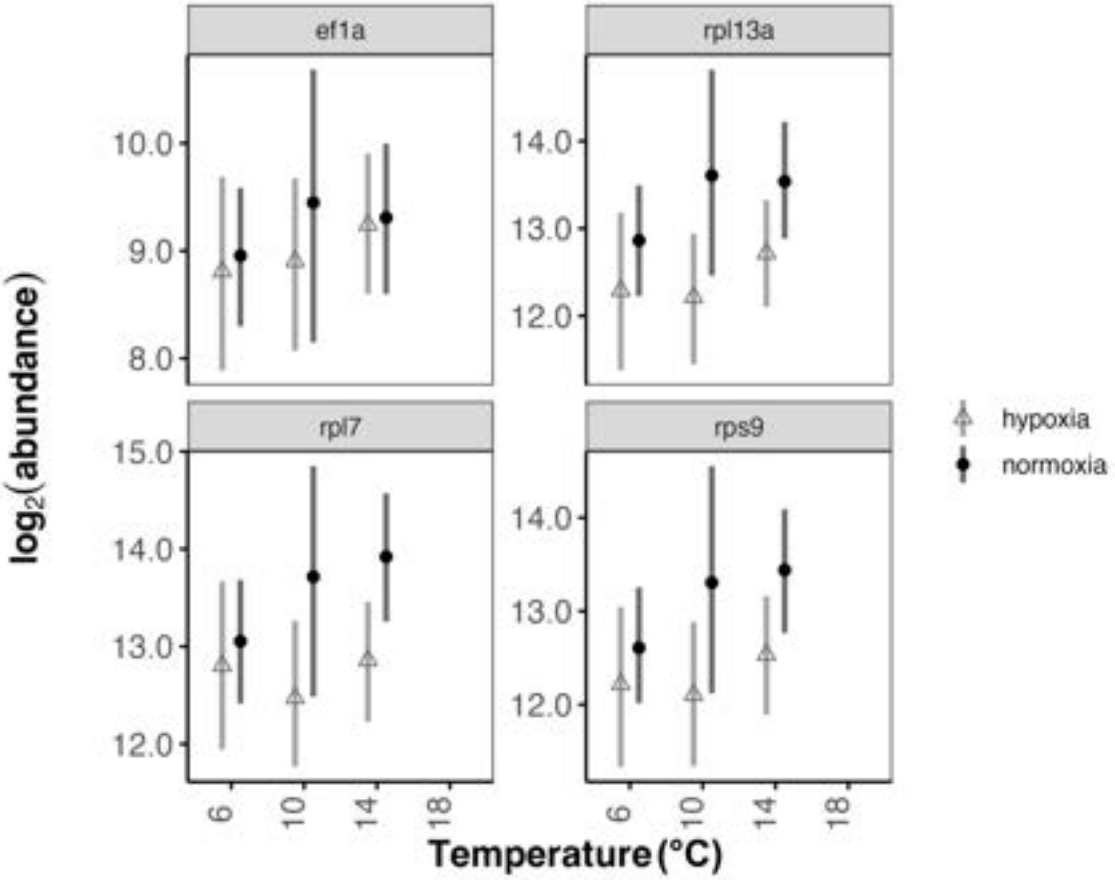


**SI Figure 12. Naïve MCMC.qpcr model of transcript abundance of housekeeping genes in the mucus across temperature acclimations and oxygen concentrations in $CT_{max}$ lake trout (*Salvelinus namaycush*).** All four housekeeping genes were used as priors in the informed model. Values from lake trout ($n$ = 5-10) acclimated to three temperatures (6, 10, 14°C) and two oxygen concentrations (normoxia: 9.93 ± 1.23 mg $L^{-1}$; hypoxia: 6.18 ± 0.44 mg $L^{-1}$).

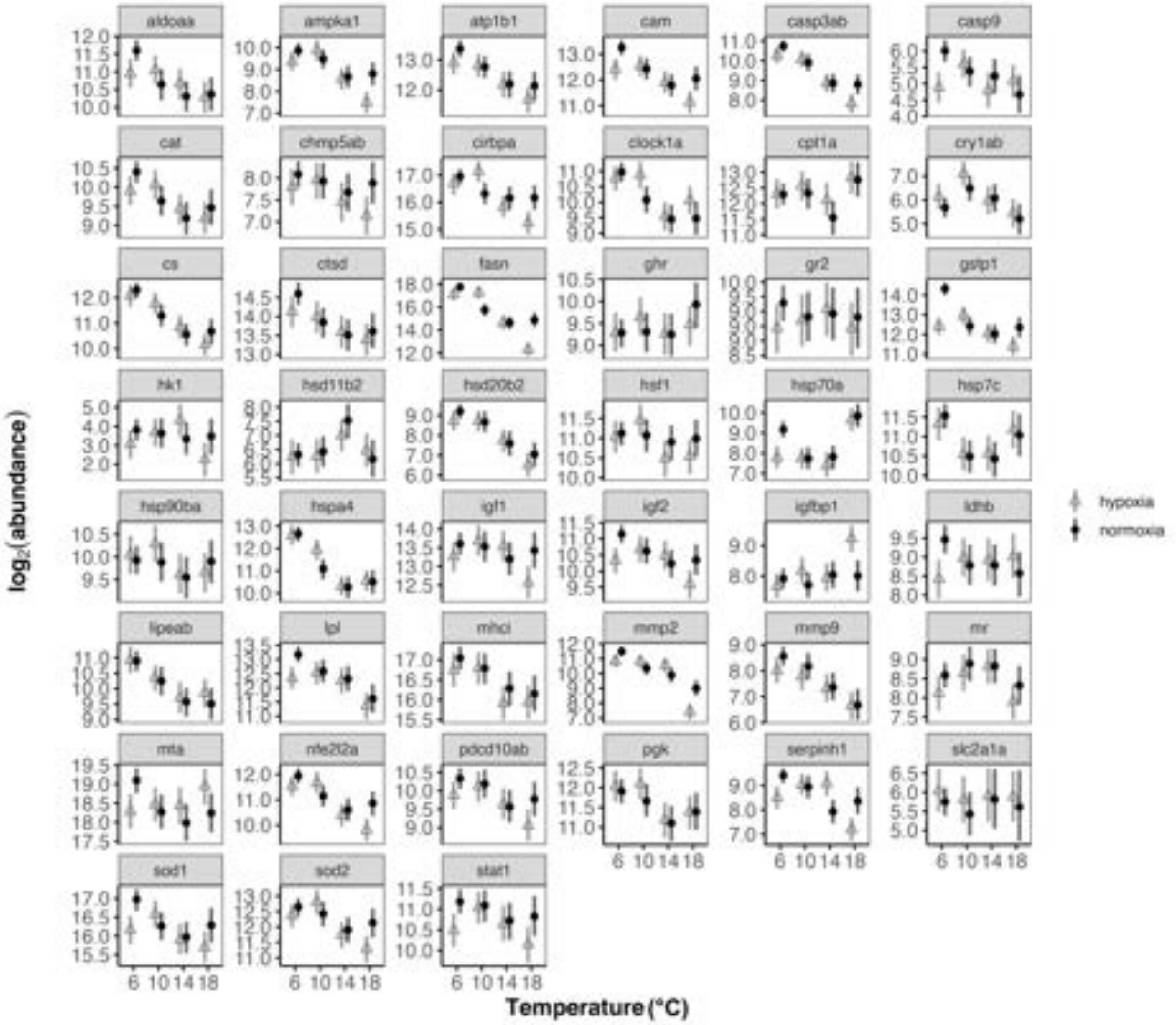


**SI Figure 13. Informed MCMC.qpcr model of all genes measured in the liver of lake trout (*Salvelinus namaycush*).** Forty-nine genes were assayed across four acclimation temperatures (6, 10, 14, 18°C) and two oxygen concentrations (normoxia: 9.93 ± 1.23 mg $L^{-1}$; hypoxia: 6.18 ± 0.44 mg $L^{-1}$) with *rpl7* used as a prior in the informed model. All housekeeping genes have been removed from this graphic and are presented separately in SI Fig. 1. Values are obtained from $n$ = 6-15 fish per treatment.

SI: Cumulative effects of high temperature and low dissolved oxygen alter the acute thermal tolerance and cellular stress response in lake trout

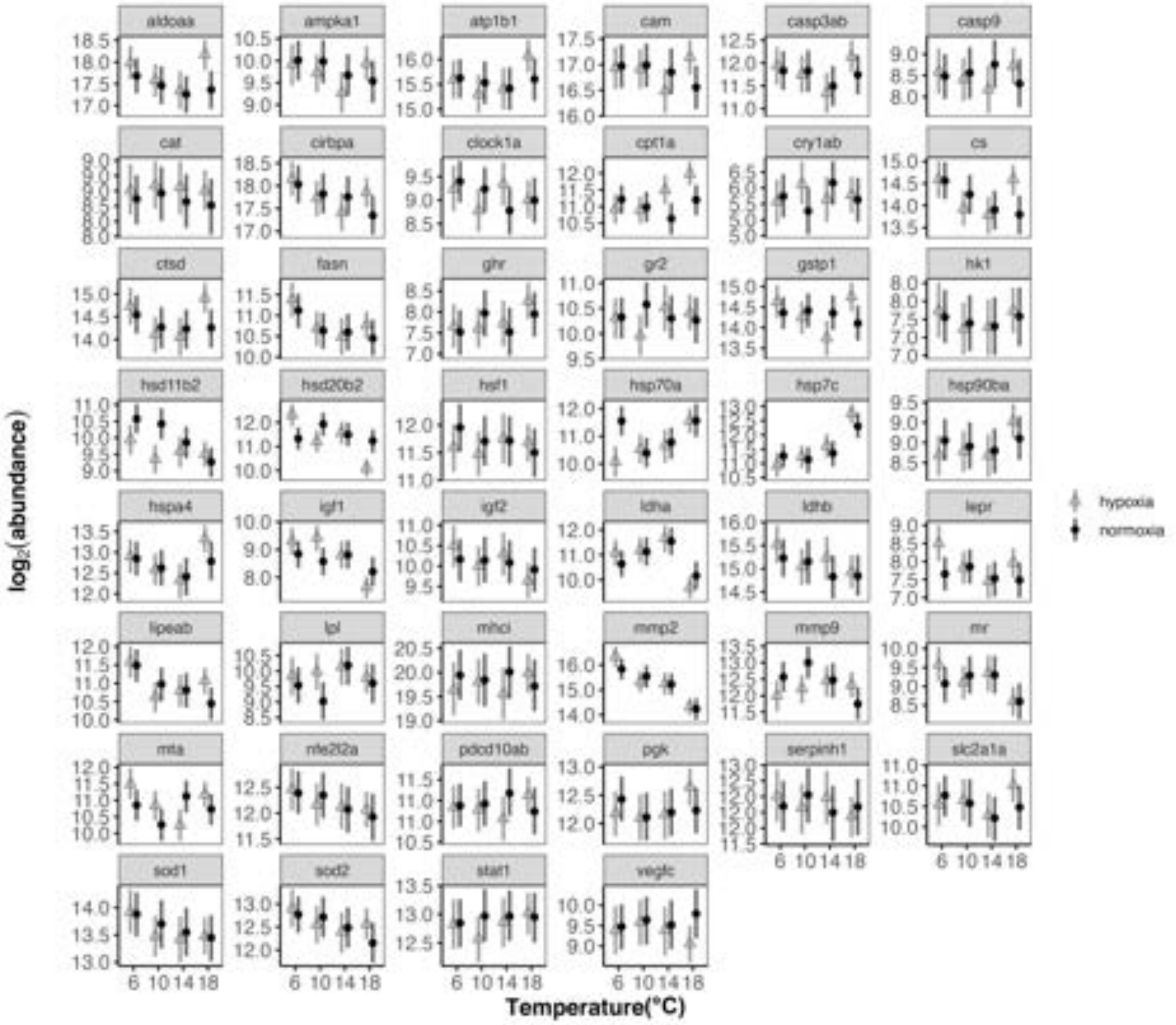


**SI Figure 14. Informed MCMC.qpcr model of all genes measured in the gill of lake trout (*Salvelinus namaycush*).** Fifty genes were assayed across four acclimation temperatures (6, 10, 14, 18 °C) and two oxygen concentrations (normoxia: 9.93 ± 1.23 mg $L^{-1}$; hypoxia: 6.18 ± 0.44 mg $L^{-1}$) with *rpl7, rpl13a,* and *rps9* were used as a prior in the informed model. All housekeeping genes have been removed from this graphic and are presented separately in SI Fig. 2. Values are obtained from $n$ = 8-13 fish per treatment.

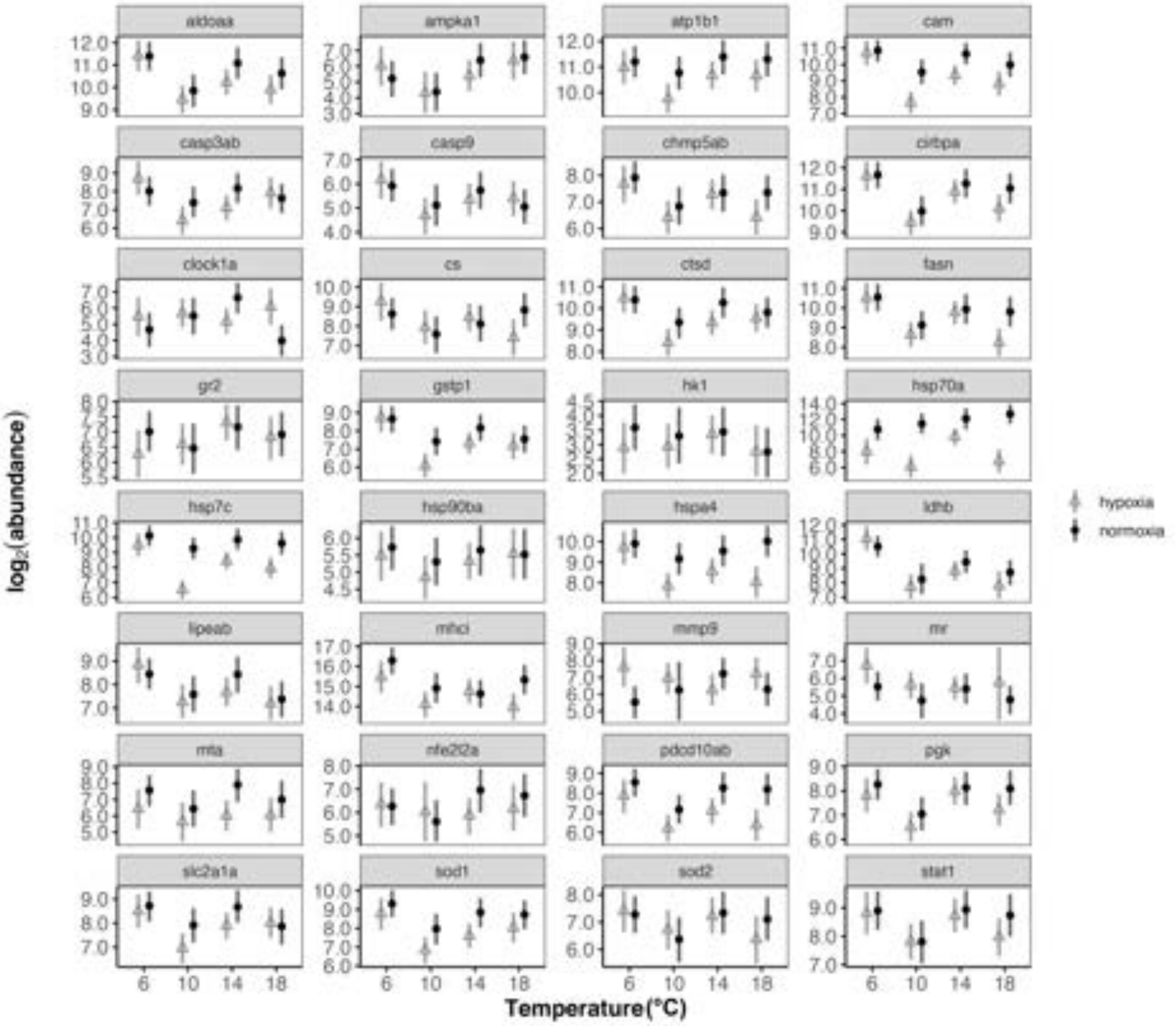


**SI Figure 15. Informed MCMC.qpcr model of all genes measured in the epidermal mucus of lake trout (*Salvelinus namaycush*).** Thirty-two genes were assayed across four acclimation temperatures (6, 10, 14, 18 °C) and two oxygen concentrations (normoxia: 9.93 ± 1.23 mg $L^{-1}$; hypoxia: 6.18 ± 0.44 mg $L^{-1}$) with *rpl7, rpl13a, rps9, ef1a* were used as a prior in the informed model. All housekeeping genes have been removed from this graphic and are presented separately in SI Fig. 3. Values are obtained from *n* = 4-9 fish per treatment.

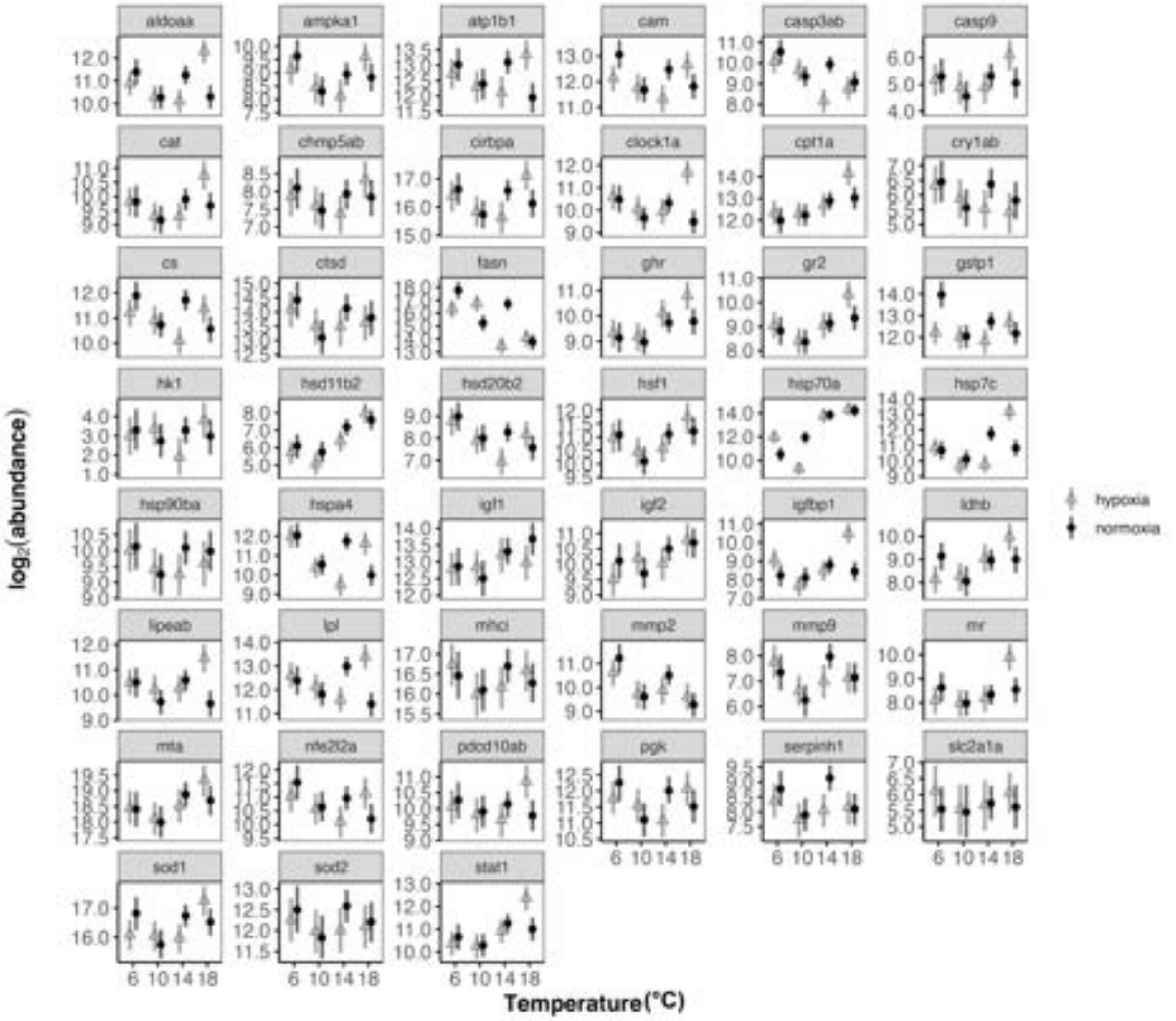


**SI Figure 16. Informed MCMC.qpcr model of all genes measured in the lake trout (*Salvelinus namaycush*) liver after CT$_{max}$.** Forty-nine genes were assayed across four acclimation temperatures (6, 10, 14, 18 °C) and two oxygen concentrations (normoxia: 9.93 ± 1.23 mg L$^{-1}$; hypoxia: 6.18 ± 0.44 mg L$^{-1}$) with *rpl7* used as a prior in the informed model. All housekeeping genes have been removed from this graphic and are presented separately in SI Fig. 10. Values are obtained from *n* = 6-12 fish per treatment.

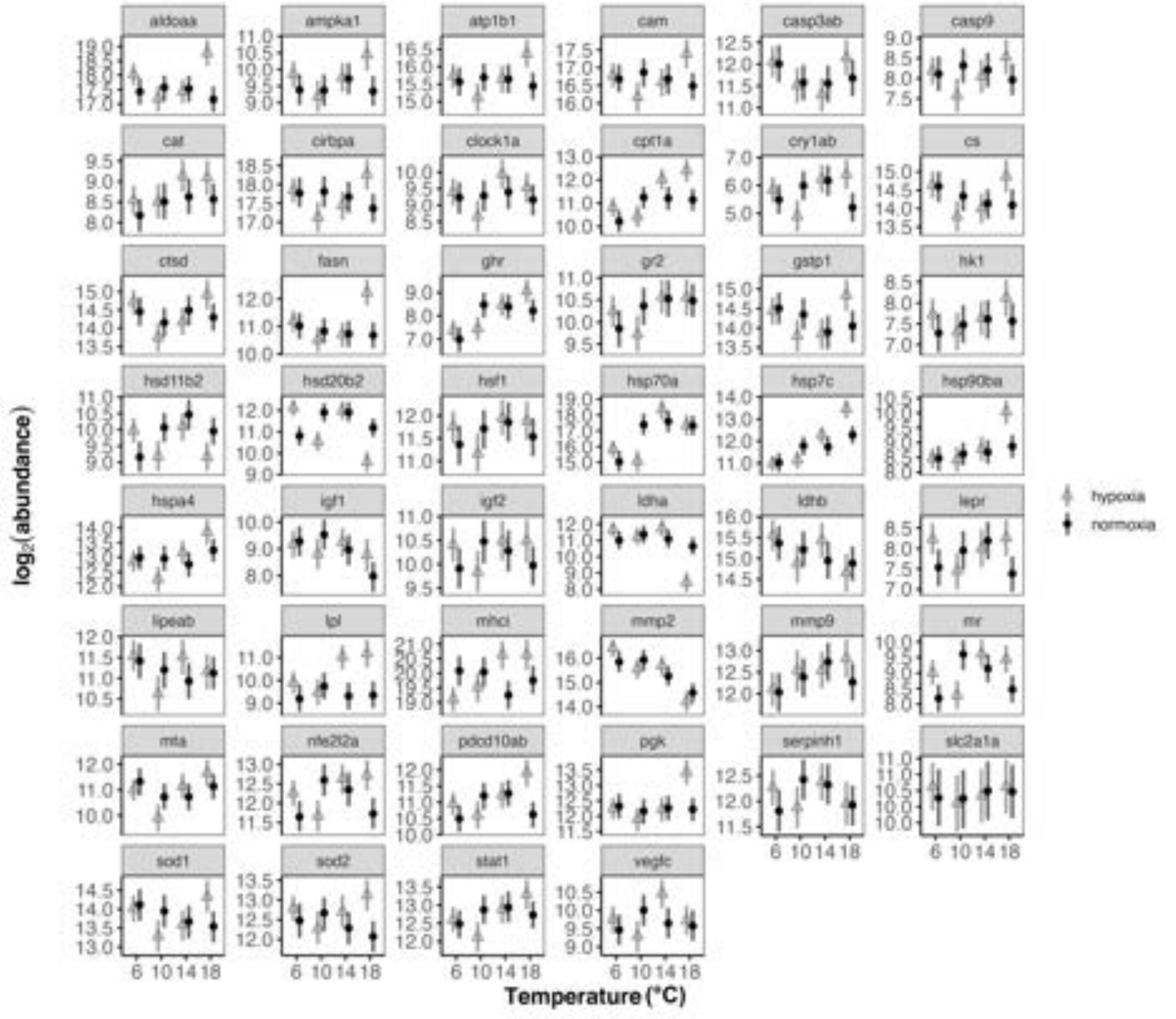


**SI Figure 17. Informed MCMC.qpcr model of all genes measured in the lake trout (*Salvelinus namaycush*) gill after $CT_{max}$.** Forty-nine genes were assayed across four acclimation temperatures (6, 10, 14, 18 °C) and two oxygen concentrations (normoxia: 9.93 ± 1.23 mg $L^{-1}$; hypoxia: 6.18 ± 0.44 mg $L^{-1}$) with *rpl7* used as a prior in the informed model. All housekeeping genes have been removed from this graphic and are presented separately in SI Fig. 11. Values are obtained from $n$ = 8-11 fish per treatment.

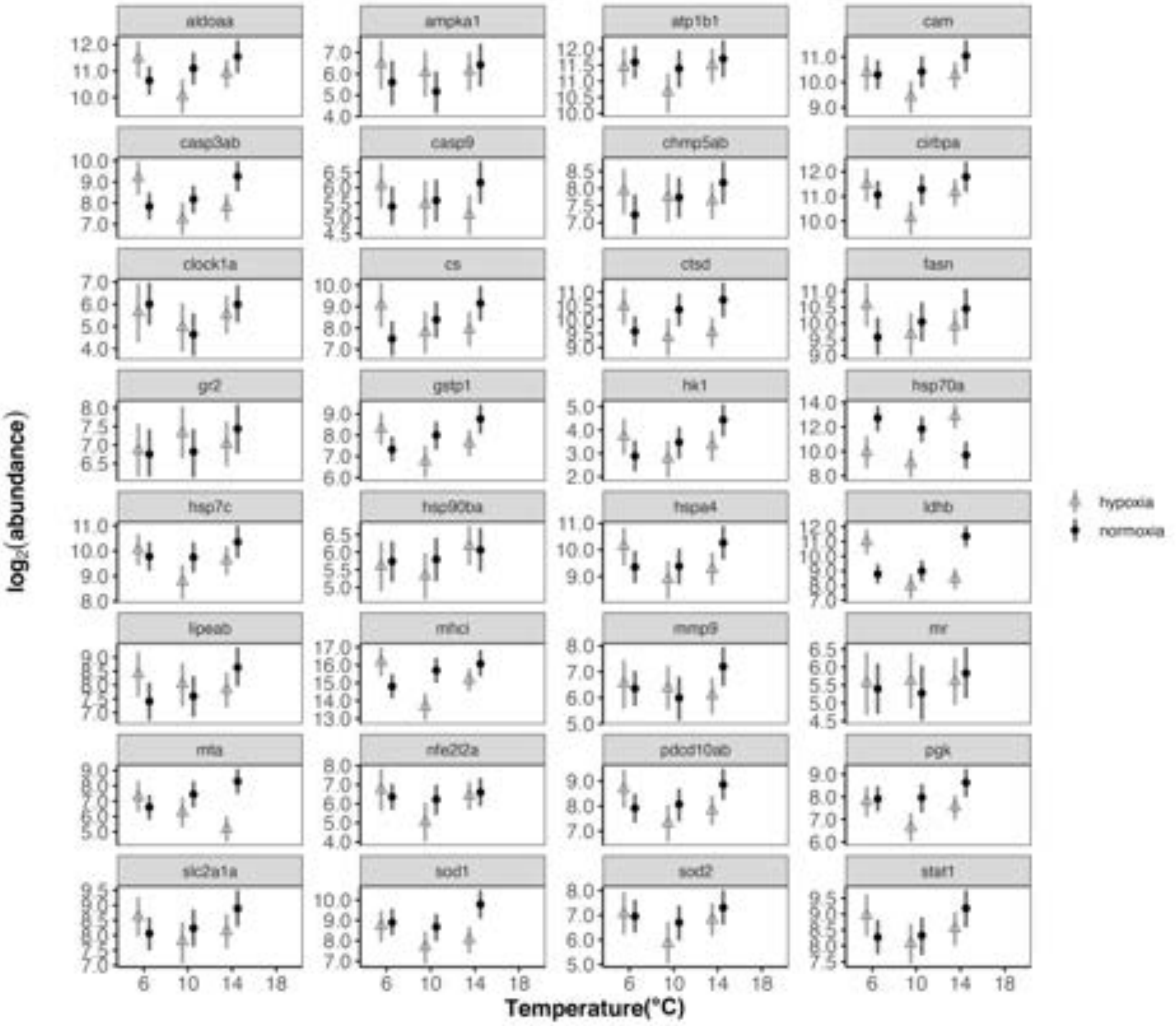


**SI Figure 18. Informed MCMC.qpcr model of all genes measured in the lake trout (*Salvelinus namaycush*) epidermal mucus after $CT_{max}$.** Forty-nine genes were assayed across three acclimation temperatures (6, 10, 14 °C) and two oxygen concentrations (normoxia: 9.93 ± 1.23 mg $L^{-1}$; hypoxia: 6.18 ± 0.44 mg $L^{-1}$) with *rpl7, rpl13a, rps9, ef1a* used as a prior in the informed model. All housekeeping genes have been removed from this graphic and are presented separately in SI Fig. 12. Values are obtained from *n* = 5-10 fish per treatment.
